# Natural non-coding *pumilio* variants retune value-coding interneurons to bias *Drosophila* oviposition choices

**DOI:** 10.1101/2025.03.07.641729

**Authors:** Dorsa Motevalli, Robert Alfredson, Sydney Fogleman, Emmanuel Medrano, Yang Chen, William I. Silander, Toshihide Hige, Ulrich Stern, Chung-Hui Yang

**Affiliations:** Dept. of Neurobiology, Duke University Medical School, Durham, NC 27710; Dept. of Biology, University of North Carolina, Chapel Hill, NC 27599; Dept. of Cell Biology and Physiology, University of North Carolina, Chapel Hill, NC 27599; Integrative Program for Biological and Genome Science, University of North Carolina, Chapel Hill, NC 27599

## Abstract

How natural regulatory genetic variation shapes innate economic decision biases by modifying neural circuit structure and function remains poorly understood. Here, we trace this pathway using a value-based oviposition decision in *Drosophila*. While laboratory flies reject sucrose in favor of a plain substrate, a wild-caught African strain accepts sucrose. This behavioral divergence maps to three African-specific intronic SNPs in the gene *pumilio* (*pum*), encoding an RNA-binding translational repressor. These SNPs downregulate *pum*, derepressing its target – the voltage-gated sodium channel *paralytic* (*para*) – in a pair of GABAergic interneurons that encode option values. Increased *para* enhances excitability, compresses neuronal value-coding differences between sucrose and plain options, and promotes sucrose acceptance. Selectively reducing *pum* or overexpressing *para* in these neurons converts laboratory flies to the African phenotype at physiological and behavioral levels. Our findings provide a genome-to-circuit-to-behavior model, illustrating how subtle regulatory polymorphisms reshape neural computations to drive adaptive variation in economic decision-making.

## Introduction

Natural genetic variation offers a powerful window into how genes fine tune neural circuits to produce behavior diversity. Single-gene variants, for example, modulate pair-bonding in voles, feeding behavior in worms, sugar-bait avoidance in cockroaches, and more recently photoperiod adaptation in *Drosophila* (*1–4*). Yet we know little about how genetic variation modifies the neural circuits that underlie value-based economic decisions – the process of ranking alternatives by their subjective worth. Ventral pallidum, orbitofrontal cortex, and dopaminergic (DA) neurons have been implicated to assign subjective option value in mammals (*5–7*), but how genetically encoded circuit differences give rise to innate decision biases remains largely unexplored.

*Drosophila* egg-laying site selection has emerged as a powerful model for studying value-based decisions (*8–12*). Female flies search, evaluate, and compare options before depositing each egg (*9, 11*). Importantly, they are capable of “choosing the greater of two goods”, a hallmark of value-based decisions. Standard laboratory strains such as *w^1118^* readily accept either a sucrose substrate or a plain substrate when only one type is offered, but they reject sucrose if both are available (**Fig. 1A**) (*9–11, 13*). Recent studies have identified the egg-laying command neurons – oviposition descending neurons (oviDNs) – in the brain (*13*). Importantly, oviDNs activity exhibits a “ramp-to-threshold” pattern (*12*) as females deliberate their choices, reminiscent of the “drift-to-bound” dynamics of decision-making neurons in higher organisms (*14, 15*). These properties, together with the extensive genetic- and circuit-manipulation tools and the recently completed connectome of an adult female brain (*16–18*), make *Drosophila* egg-laying decisions ideal to explore how genetically encoded circuit variations drive variations in decision outcomes in nature.

**Fig. 1.**
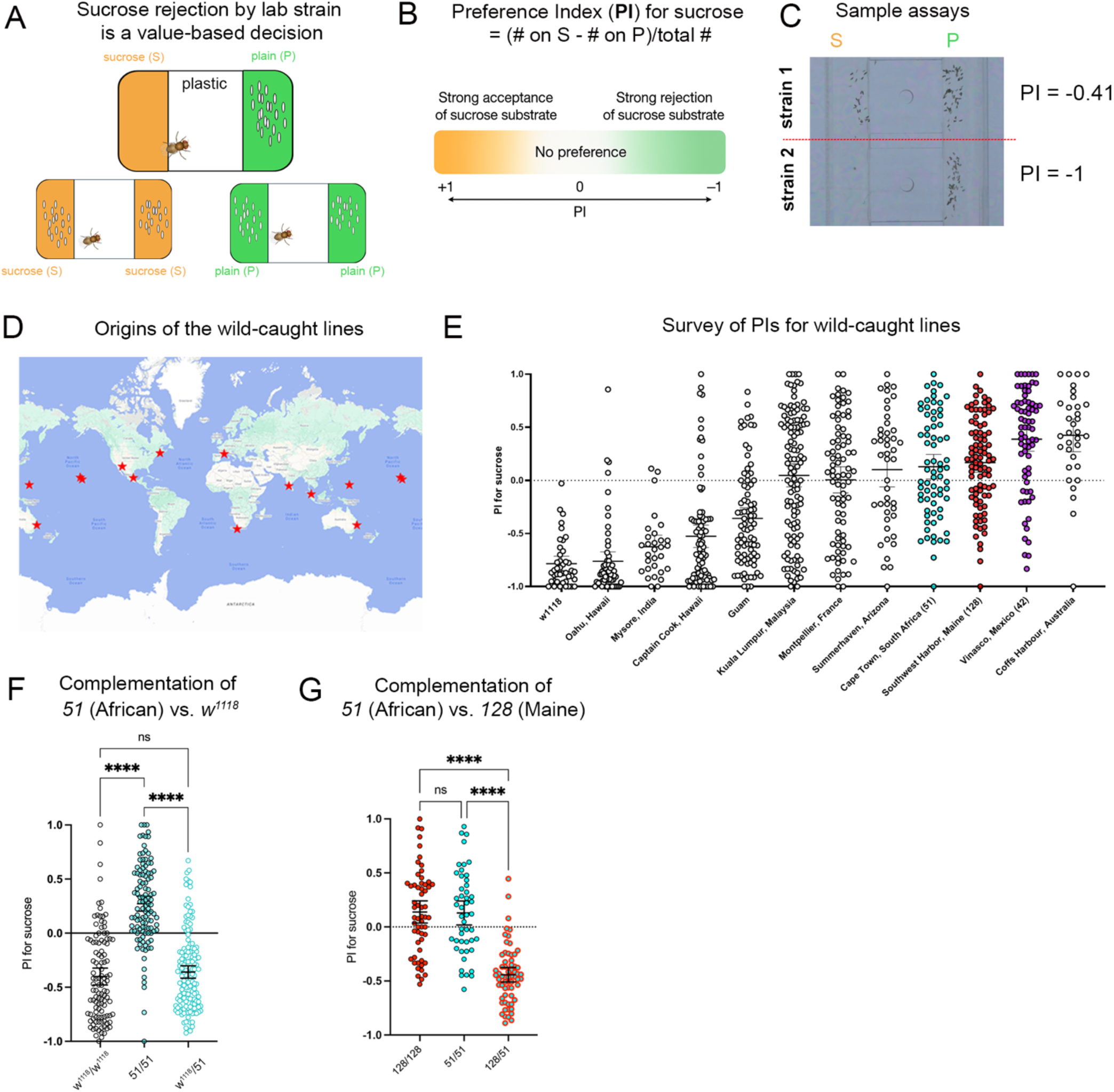
Genetic control of natural = variation in sucrose preference during oviposition decision. **(A)** Schematics illustrating that decision between a sucrose (150 mM sucrose in 1% agarose) and a plain (1% agarose) substrate is value-based: standard lab strain *w^1118^* accepts either substrate when offered alone but rejects sucrose in a choice test. **(B)** Definition of the Preference Index (PI) for sucrose (PI = [eggs on S – eggs on P] / total eggs). PI = +1 indicates full sucrose preference, –1 full rejection, 0 no bias. **(C)** Representative choices from two individual females (see also **fig. S1** for tool for automatic counting). **(D)** Geographic origins of the wild-caught lines tested. **(E)** PI distribution across strains; each point represents PI of one fly. **(F)** PIs of parental strains *w^1118^* and line 51 and their F₁ hybrid (*w^1118^/51*); n = 111, 113, 149. **(G)** PIs of parental strains *128* and 51 and their F₁ hybrid (*128/51*); n = 58, 48, 64. Homozygous lines are denoted “*51/51*” or “*128/128*” (as opposed to 51 and 128) at times when emphasis is needed. Unless noted, statistics and full sample-size information are provided in Material and Methods.

Here, we exploit a striking difference in sugar preference during egg-laying between the standard laboratory strain *w^1118^* and a wild-caught African strain to investigate how natural genetic variation sculpts the circuitry to generate distinct innate decision biases. By combining behavior analysis, genome-wide SNP mapping, and connectome-guided circuit interrogation, we pinpoint differential expression of *pumilio* (*pum*), encoding an evolutionarily conserved RNA-binding protein (*19, 20*), as one key determinant governing the two strains’ distinct choices between sucrose and plain substrates. Strain-specific intronic SNPs tune *pum* expression, which in turn adjusts the level of voltage-gated sodium channel encoded by *paralytic* (*para*), a known target of *pum* (*21*). This *pum*-adjusted difference in excitability alters the ratio of the sucrose-versus plain-evoked response in a pair of value-coding – and decision-critical – inhibitory interneurons we identified, thereby biasing the animal’s behavioral choice. Together, these results demonstrate that genetically encoded differences in *pum* expression can modify circuit properties, offering a rare end-to-end example of how natural variation in a gene-regulatory region reshapes computations within a defined microcircuit to yield strain-specific decision outcomes that may better suit each habitat.

## Results

### Sucrose preference varies among wild-caught strains and is under simple genetic control

To investigate whether strains originated from different locations in the world display different innate oviposition decision biases, we surveyed a collection of wild-caught lines from the Cornell Stock Center for their sucrose-versus-plain choices (**Fig.1A-E**). We used custom apparatuses and egg-counting code to assay preferences of many animals in parallel (*22*) (**Extended Data Fig.1**) and calculated each individual’s sucrose preference index (PI) as a quantitative measure of decision bias (**Fig.1B, C**). As anticipated, not all wild-caught strains rejected sucrose; while our standard laboratory strain (*w^1118^*) rejected sucrose and had a mean PI of ∼ - 0.7, many of the wild-caught strains were either indifferent towards – or even preferred sucrose – for egg-laying (**Fig. 1E**).

To probe the genetic complexity underlying sucrose acceptance differences, we performed complementation to test how the F_1_ progeny of the standard lab strain *w^1118^* and the sucrose-accepting African strain line 51 would choose. Interestingly, these F_1_ progeny (*w^1118^/51*) behaved like *w^1118^* and rejected sucrose (**Fig. 1F**), suggesting sucrose acceptance is a recessive trait. Further, the F_1_ progeny (*128/51*) of two sucrose-accepting strains (lines 51 and line 128 from Maine) rejected sucrose (**Fig. 1G & Extended Data Fig. 2**). This result not only confirms that sucrose acceptance is a recessive trait but also indicates that the causal variants in nature occur in at least two complementation groups – that is, they have at least two distinct genetic origins. These observations suggest that: 1) the genetic control of how animals value sucrose in this decision is relatively simple; 2) the SNPs that drive sucrose rejection versus acceptance could in principle be mapped. In this paper, we use the term SNPs in a broad sense to include both single nucleotide polymorphisms and indels (insertions and deletions).

### Intronic SNPs in the RNA-binding translational repressor pumilio (pum) in the African strain drive sucrose acceptance

To map the SNPs – and thus the genes – underlying the laboratory (*w^1118^*) and African (line 51) strains’ divergent valuation of sucrose (**Fig. 2A**), we generated animals with hybrid genomes derived from both strains. We backcrossed the F_1_ female progeny of the *w^1118^* × line 51 cross to line 51 males, so that each F_2_ female carried one non-recombinant chromosome from line 51 and one recombinant chromosome (of line 51 and *w^1118^*) (**Fig. 2B**). We then assayed the PI for sucrose of many F_2_ individuals (**Fig. 2C**) and selected 192 of them for full-genome sequencing. We next conducted a GWAS with PLINK (*23*) to identify SNPs associated with PI (**Fig. 2D**). Three statistically significant, high-impact (i.e., large effect) SNPs are particularly notable because they lie within the introns of the same gene: *pumilio* (*pum*) (**Fig. 2E & Supplementary Table 1**), which encodes an RNA-binding translational repressor (*19, 20*).

**Fig. 2.**
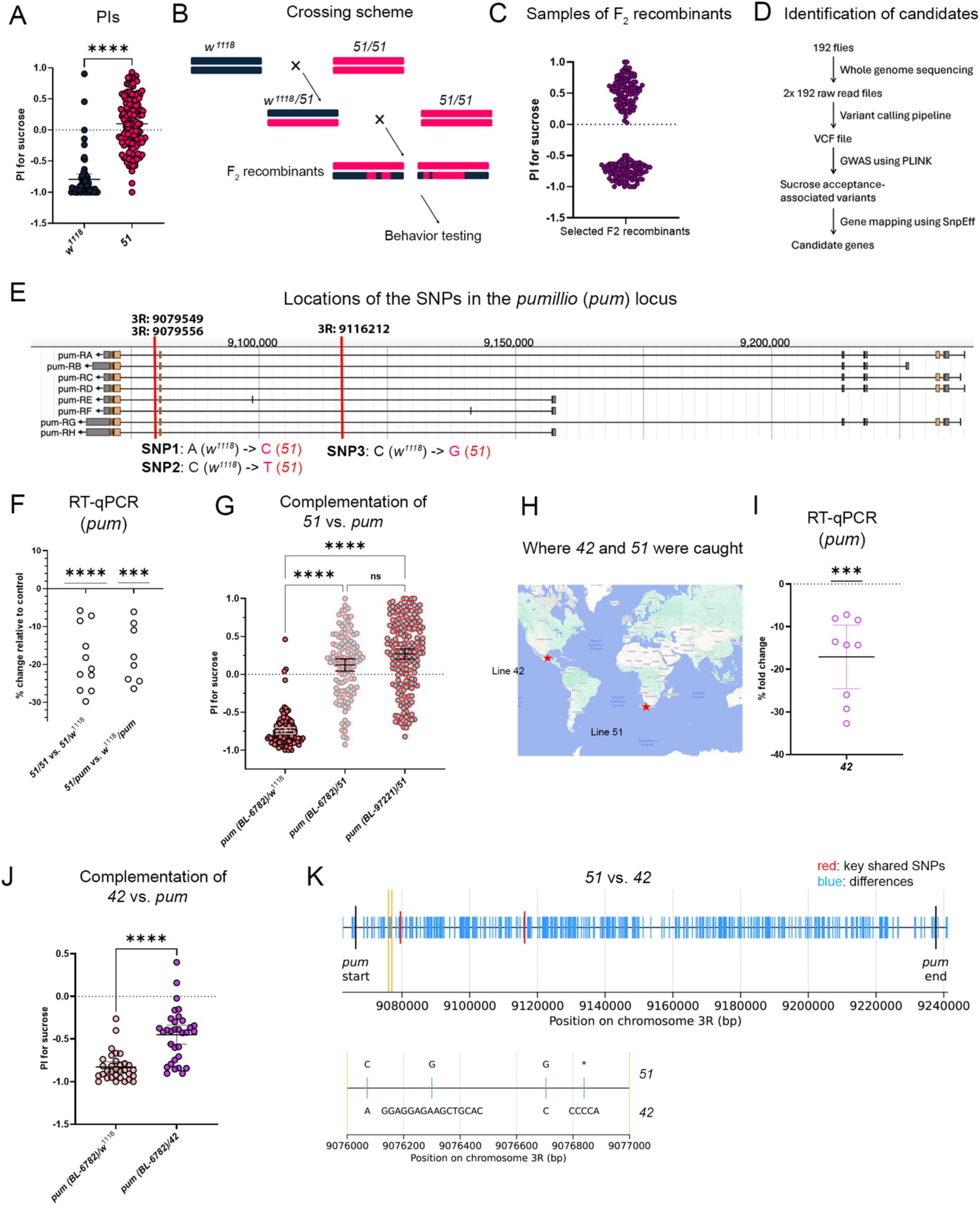
Intronic *pum* SNPs in wild-caught strains lower *pum* expression and increase sucrose acceptance. **(A)** PIs of control (*w*^1118^) vs. line 51 (n = 63, 141). **(B)** Crossing scheme to produce F₂ recombinants. **(C)** PIs of sample F₂ recombinants. **(D)** GWAS pipeline for mapping the trait. **(E)** Gene structure of *pum* with three intronic significant SNPs marked (orange). **(F)** RT-qPCR of brains showing reduced *pum* expression in *51/51* vs. *51/w^1118^* (and in 51/*pum* vs. *w*^1118^/*pum* (n = 11, 8 biological replicates). Expression normalized to *actin5C*; one-sample t-test vs. 0. **(G)** Complementation of two *pum* alleles crossed to line 51 (n = 111–182). **(H)** Origins of lines 42 (Mexico) and 51 (Africa). **(I)** RT-qPCR of heads showing reduced *pum* expression in line 42 (n = 9 biological replicates). Expression normalized to *actin5C*; one-sample t-test vs. 0. **(J)** Complementation test of a *pum* allele crossed to line 42 (n = 32, 32). **(K)** Sequence divergence between 51 and 42. Lower panel: magnified view of the region bracketed by two orange bars. * = missing bp.

Given sucrose acceptance in egg-laying decision is a recessive trait, we speculated that the SNPs in line 51 reduce *pum* expression. This is because recessive “mutations” tend to reduce gene expression/function. Indeed, *pum* expression was lower in the brains of line 51 than controls (**Fig. 2F**). We next asked whether line 51 would fail to complement *pum* mutations by assessing the F_1_ progeny of line 51 and different *pum* mutants, a direct test for a causal role of reduced *pum* in driving sucrose acceptance. While F_1_ progeny from control crosses (*pum*/*w^1118^*and *51/w^1118^*) rejected sucrose, *pum/51* accepted sucrose; importantly, this was true for two different *pum* alleles from different genetic backgrounds (**Fig. 1F & Fig. 2G**). We then asked if the SNPs we had identified in line 51 might be present in other sucrose-accepting wild-caught strains. By looking for other lines that failed to complement line 51 (**Extended Data Fig. 2**), we found that line 42 – a line caught in Mexico – indeed harbored the same SNPs (**Fig. 2H and Supplementary Table 2**). Consistent with the idea that these SNPs reduce *pum* expression, line 42 flies showed lower *pum* expression (**Fig. 2I**) and failed to complement a *pum* mutation, too (**Fig. 2J**). Notably, despite sharing the three key intronic SNPs in *pum*, these two strains are otherwise genetically divergent. They differ at many positions within the *pum* locus, including several sizeable deletion–insertion variants, reflecting their distinct geographical origins (**Fig. 2K and Supplementary Table 2**).

Together, our analyses suggest that the level of *pum* expression – regulated in part by its intronic sequences – is crucial for how *Drosophila* value sucrose relative to plain substrates during egg-laying decisions. Specifically, higher *pum* expression in the laboratory strain *w^1118^* drives sucrose rejection, whereas lower *pum* levels in the African strain line 51 (and the Mexican strain line 42) promote sucrose acceptance.

### Sucrose rejection in the lab strain is mediated by a pair of interneurons Earmuff

To pinpoint the neural substrates through which reduced *pum* expression promotes sucrose acceptance in the African strain, we first identified – in the laboratory strain – the critical circuit that converts the sensory qualities of egg-laying options into value signals relayed to the command neurons oviDNs (**Fig. 3A**). A recent study showed that leg sweet-taste neurons and their postsynaptic targets, TPN2 neurons (**Fig. 3B**), are necessary and sufficient to devalue sucrose as the laboratory strain deliberates between sucrose and plain substrates (*22*) (**Extended Data Fig. 3**). In addition, a few recurrently connected excitatory neurons (e.g., oviENs) that signal oviDNs have been reported (*12, 13*). However, the circuit components that bridge the TPN2 axons – which terminate in the subesophageal zone (SEZ) – and the recurrently connected groups presynaptic to the oviDNs – which arborize in the superior lateral protocerebrum (SLP) – remain unknown (**Fig. 3C**). We performed a small-scale screen where we silenced some neurons projecting from the SEZ to the SLP with Kir channel, using the morphological criteria and *split-GAL4* lines described by Sterne et al. (*24*) (**Fig. 3D**). Inactivating neurons labeled by one specific line consistently increased sucrose acceptance (**Fig. 3E, F**). These neurons are called Earmuff likely because their morphology resembles a pair of earmuffs (**Fig. 3G & Fig. 4A**). (Note that all the transgenic tools we used for this work were generated in the genetic background of the standard lab strain, which rejects sucrose when assayed individually.)

**Fig. 3.**
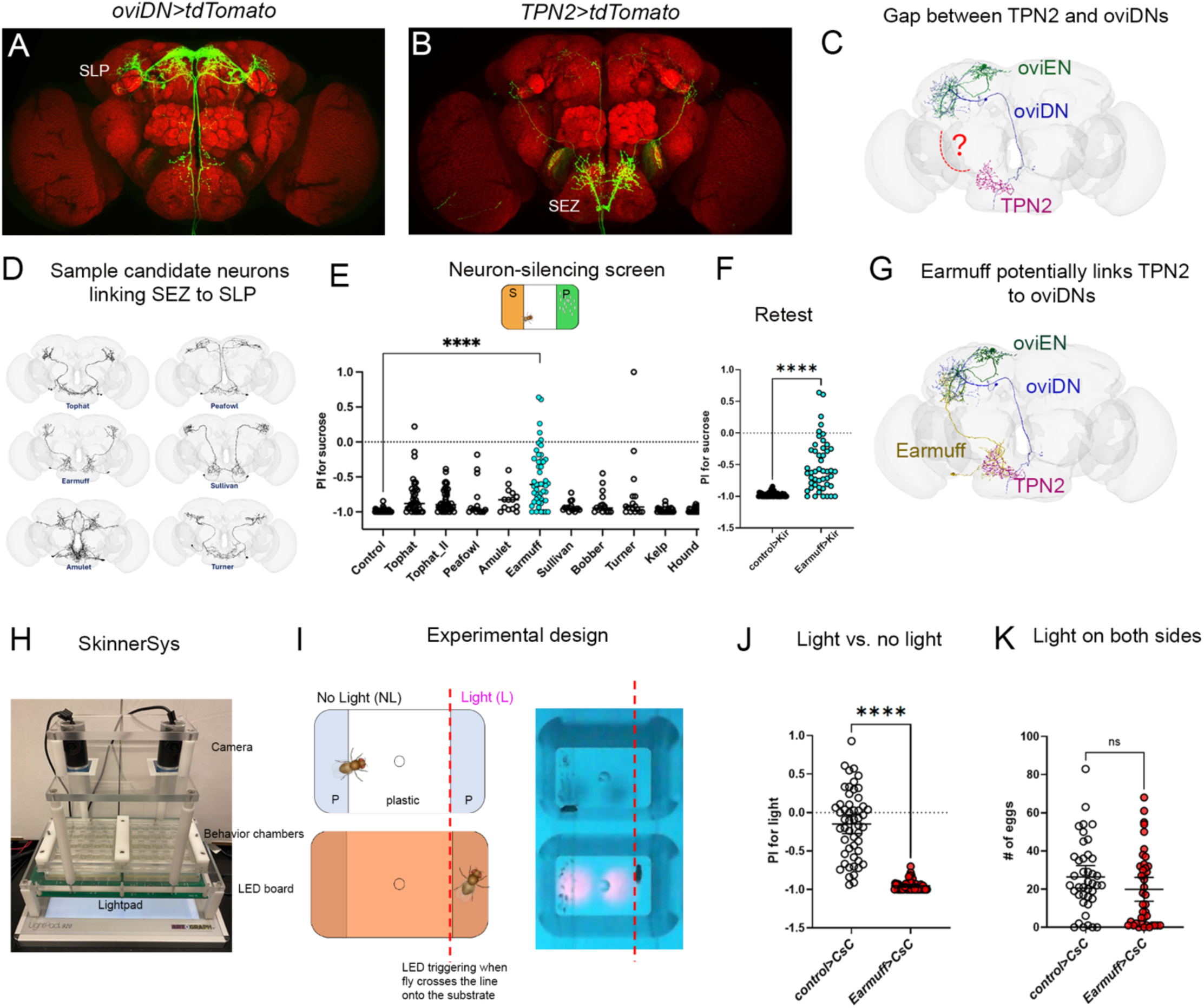
Earmuff neurons are necessary and sufficient for sucrose rejection in the lab strain (*w^1118^*). **(A, B)** False-color confocal images of oviDN neurons and TPN2 (cell bodies in VNC; see **fig. S3**). **(C)** FlyWire reconstructions of TPN2, oviEN, and oviDN in one hemisphere; dashed red line indicates the missing component(s) in the TPN2 → oviEN/DN pathway. **(D)** Sample SEZ → SLP projection neurons we screened. **(E)** PIs of flies with Kir-mediated silencing of candidate neurons. **(F)** Independent validation of silencing Earmuff neurons. **(G)** FlyWire reconstructed morphology of TPN2, Earmuff, oviEN, and oviDN in one hemisphere. **(H)** SkinnerSys rig, which delivers real-time optogenetic stimulation to up to 40 flies. A camera tracks behavior in individual chambers positioned above an LED array. **(I)** Stimulation paradigm: when a fly remains on the non-light (NL) side (left of the pink midline) no light is delivered; crossing to the light (L) side triggers illumination. Right: Representative camera frames during stimulation. LEDs appear dim because of a filter we placed in front of the cameras. **(J)** PIs of *Earmuff>CsChrimson* flies vs. controls in a light/no-light choice (n = 52, 54). **(K)** Total eggs laid with light triggered on both substrates (n = 41, 40; p = 0.13). Note that all the transgenic lines we used for imaging and manipulating neurons in this work were generated in the background of the lab strain (*w^1118^*).

**Fig. 4.**
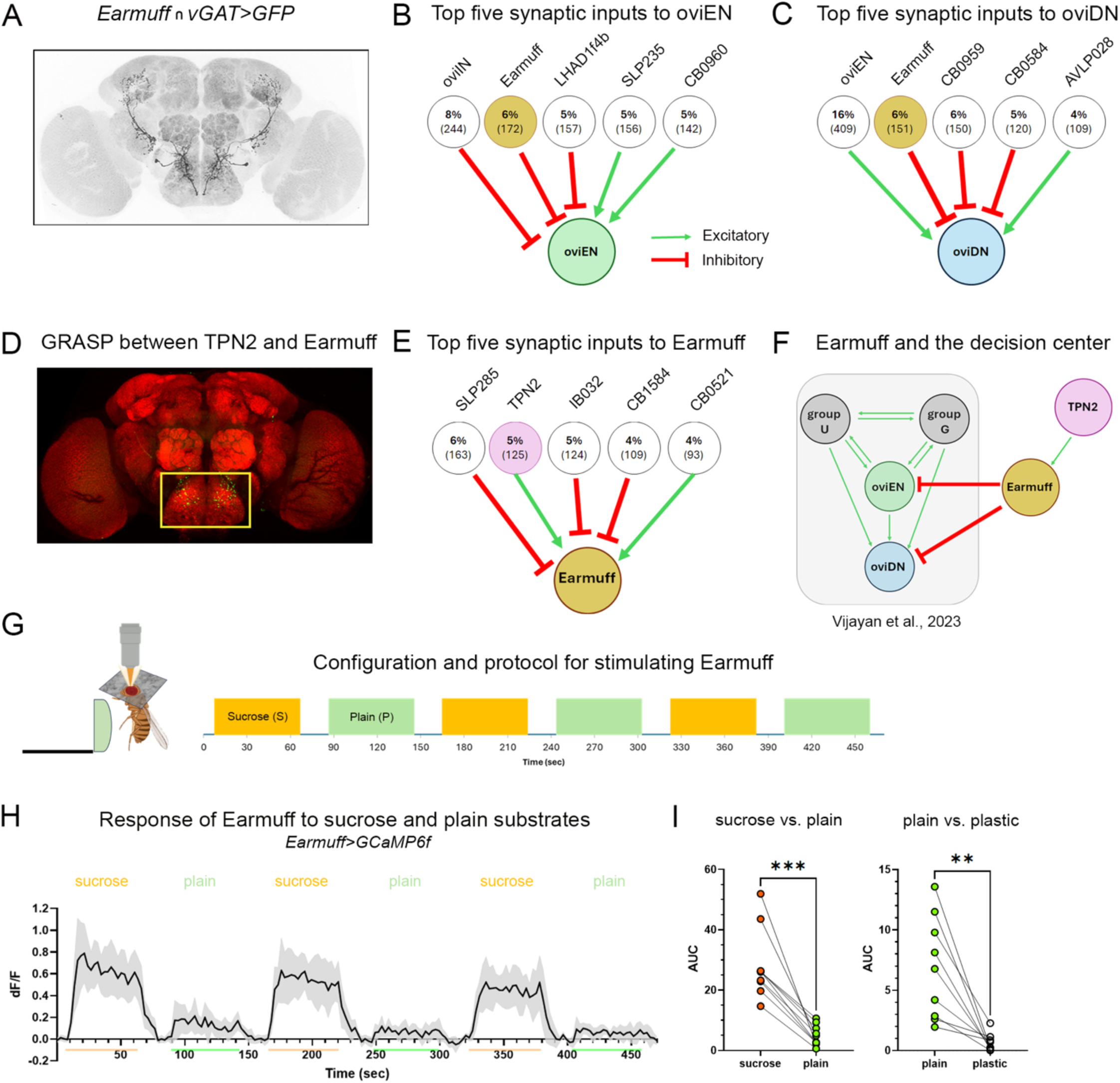
Earmuff neurons prefer sucrose in the lab strain (*w^1118^*) and inhibit the decision node. **(A)** *vGAT-lexA* is active in Earmuff, indicating these neurons are GABAergic. **(B, C)** Top five presynaptic partners of oviEN and oviDN according to FlyWire. **(D)** GRASP signal (green puncta in the yellow box) marks TPN2 → Earmuff anatomical contacts in the SEZ. **(E)** Top five presynaptic partners of Earmuff according to FlyWire. **(F)** A model illustrating the circuit relationships among the command neurons *ovi*DNs, their behaviorally validated excitatory inputs, and the inhibitory Earmuff neurons. Green arrows: excitatory connections; red lines: inhibitory connections. The gray-shaded portion is adapted from Vijayan et al., 2023 (*12*). **(G)** Configuration and protocol for imaging substrate responses of Earmuff. **(H)** Mean ΔF/F of Earmuff neurons to sucrose (orange bar) and plain (green bar) (n = 10). Shaded areas: 95% CIs. **(I)** Left, AUCs for sucrose vs. plain (n = 10); right, AUCs for plain vs. plastic (n = 9). AUC used because Earmuff responses tend to be sustained.

Having found that Earmuff neurons are necessary for proper sucrose rejection, we asked if their artificial activation is sufficient to promote rejection. We expressed the channel-rhodopsin *CsChrimson* (*25*) in them and assessed how flies chose between two plain substrates, only one of which triggers light upon contact (**Fig. 3H-K**). We implemented this using SkinnerSys, a system we developed that can track 40 animals in parallel (per apparatus) and deliver light to them in closed-loop (*22*) (**Fig. 3H, I**). *Earmuff>CsChrimson* (*CsC*) animals robustly rejected the illuminated plain option whereas controls flies were indifferent between the two options (**Fig. 3J**). To determine whether optogenetic activation of Earmuff neurons simply shuts down egg-laying or “devalues” the option on which it occurs, we assessed how the flies behaved when both plain options were illuminated (**Fig. 3K**). Flies did not reduce their egg-laying rate significantly when their Earmuff neurons were optogenetically activated on both options (**Fig. 2F**), suggesting that Earmuff activation *reduces the value* of an option as opposed to just abolishing egg-laying. This mirrors how flies behave towards sucrose: they refrain from egg-laying on sucrose when a plain substrate is available but accept sucrose when it is the only option (**Fig. 1A**).

### Earmuff neurons inhibit the decision-maker and encode option values

We next investigated the circuit mechanism by which Earmuff neurons promote sucrose rejection in the lab strain. Assessing the relationship between Earmuff and the command neurons oviDNs (*16*) in the Flywire connectome revealed two noteworthy features. First, Earmuff neurons are annotated to be GABAergic, and we confirmed this by showing that the promoter for *vGAT*, a marker for GABAergic neurons, was active in them (**Fig. 4A**). Second, Earmuff neurons are annotated to directly signal both oviENs – the key excitatory input to oviDNs (*12, 13*) – and to oviDNs themselves (**Fig. 4B, C**). In fact, Earmuff neurons provide the second-strongest inputs to each target (**Fig. 4B, C**). These, together with the behavioral phenotypes, suggest Earmuff neurons contribute to sucrose rejection by inhibiting the decision-maker (oviDNs) and its strongest excitatory drive (oviENs).

How could inhibition from Earmuff neurons enable flies to reject sucrose while accepting plain substrate? One possibility is that Earmuff neurons are activated specifically by sweet taste, so that only the sucrose option triggers their inhibition of the decision-making circuit. Ca^2+^ imaging show that Earmuff neurons indeed responded strongly to sucrose solution, “opto sugar”, and sucrose agarose substrate (**fig. S4**), and were functionally connected to sweet taste projection neuron TPN2 previously shown to be important for sugar rejection (*22*) (**Fig. 4D, E, & Extended Data Fig. 4**). Interestingly, however, we found that Earmuff neurons responded to the plain agarose substrate, too, albeit at a reduced level (**Fig. 4G, H, I**) but they showed little or no response to a hard plastic surface (**Fig. 4I**). These results suggest that Earmuff neurons are sensitive to both sweet taste and texture of an egg-laying substrate, in keeping with a recent connectome-based analysis showing that multi-modal integration is a common theme of higher-order “taste neurons” (*26*).

Our combined behavioral, anatomical, and physiological evidence thus indicates that Earmuff neurons encode innately specified, substrate-specific “negative values” for sucrose and plain options, enabling the decision-maker to distinguish between them. In the lab strain, sucrose substrate activates Earmuff neurons strongly, leading to greater inhibition of the decision-maker and a lowered perceived value. Conversely, the plain substrate activates Earmuff neurons mildly, inhibiting the decision-maker less and therefore carrying a higher value than sucrose. Curiously, while males lack oviEN/DN neurons, their Earmuff neurons are morphologically indistinguishable from those of females and are likewise anatomically connected to TPN2 neurons (**Extended Data Fig. 5**).

### pum expression level drives strain-specific Earmuff value coding and decision outcomes

Having found that differential responses of Earmuff neurons to sucrose and plain substrates are likely critical for sucrose rejection over plain in the lab strain, we next asked whether sucrose acceptance in the African strain results from altered substrate responses of its Earmuff neurons (**Fig. 5A**). To assess how Earmuff neurons respond to substrates in the African strain, we used the trans-heterozygous *51/pum* flies as surrogates. We initially expected Earmuff neurons in *51/pum* animals would have reduced response to sucrose substrates. To our surprise, sucrose responses of Earmuff in these animals were comparable to controls, but their responses to plain increased significantly, elevating the response ratio of plain to sucrose (**Fig. 5B & Extended Data Fig. 6**). We also examined responses of Earmuff in trans-heterozygous *42/pum* flies (the surrogates for the Mexican line) and found a similar phenotype (**Fig. 5C**). These results suggest Earmuff’s ability to discriminate between sucrose and plain substrates is diminished in both sucrose-accepting wild-caught strains; the underlying cause, however, is an increased response to the plain substrate.

**Fig. 5.**
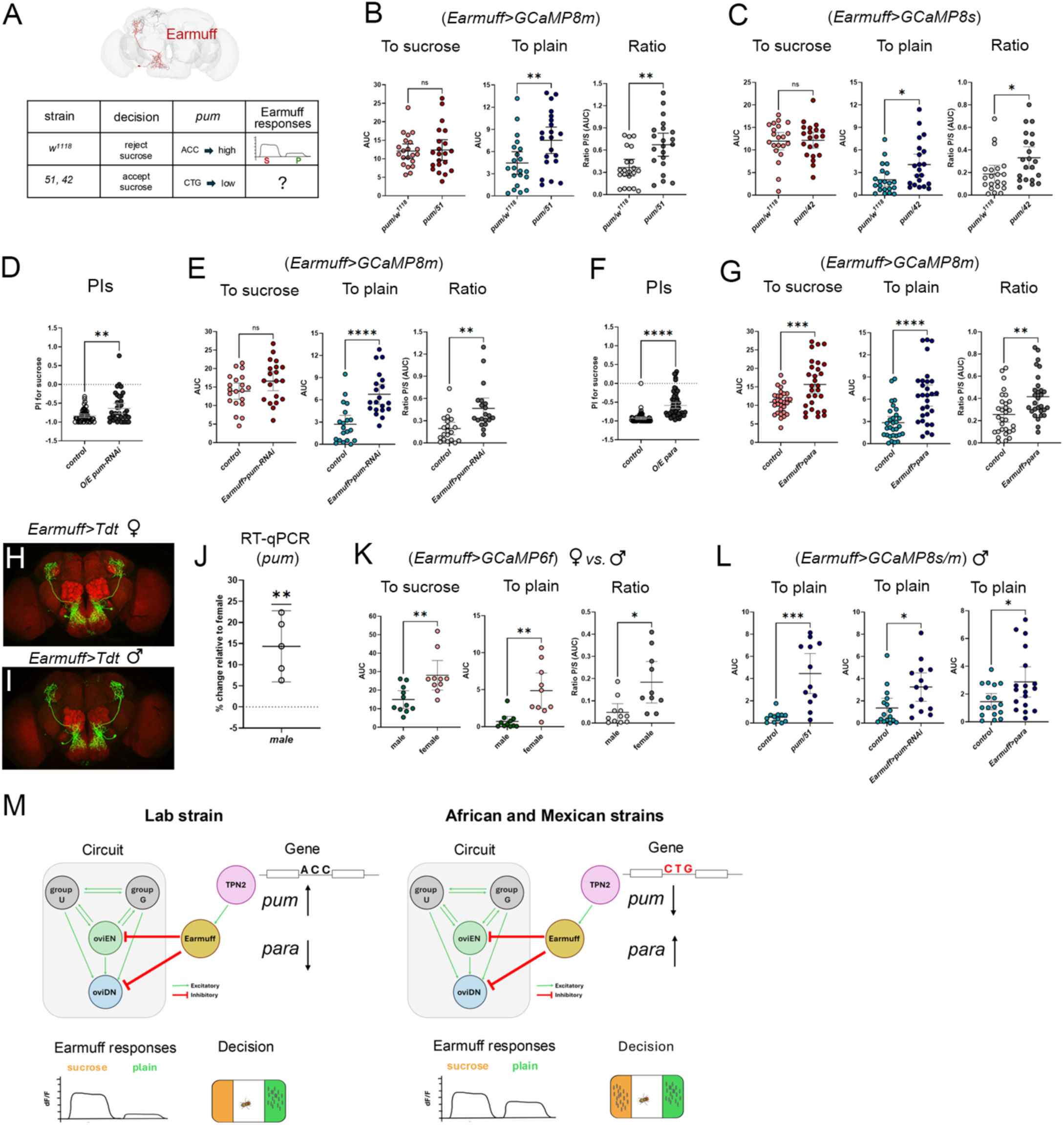
*pum* levels in Earmuff neurons set strain-specific value coding and sucrose preference. **(A)** Table summarizing results we have presented so far. **(B)** Earmuff responses in *pum*/*w^1118^*(control) vs. *pum/51* (African surrogate); n = 21 each. Left to right: AUC sucrose, AUC plain, and plain/sucrose ratio. **(C)** Earmuff responses in *pum*/*w^1118^* vs. *pum/42* (Mexican surrogate); n = 21 each. **(D)** PIs after Earmuff-specific *pum* knockdown in *w^1118^* (n = 54, 36). **(E)** Earmuff responses after Earmuff-specific *pum* knockdown in *w^1118^* (n = 20 each). **(F)** PIs after Earmuff-specific *para* overexpression in *w^1118^*(n = 72, 65). **(G)** Earmuff responses after Earmuff-specific *para* overexpression in *w^1118^* (n = 30, 29). **(H**, **I)** Earmuff neurons in both sexes. **(J)** *pum* level in males against females in *w^1118^* (n = 5 biological replicates; normalized to *actin5C*). **(K)** Earmuff responses in females vs. males in *w^1118^* (n= 10, 11). **(L)** Male Earmuff responses in *51/pum* (n = 12, 12), after Earmuff-specific *pum* knockdown (n = 16, 14), after Earmuff-specific *para* overexpression (n = 17, 18) in *w^1118^*. **(M)** Our model. In *w^1118^*, intronic SNPs elevate *pum* expression in Earmuff, lowering their excitability, increasing the contrast the sucrose-verus-plain code, and driving sucrose rejection. In wild-caught strains, corresponding SNPs reduce *pum*, increase excitability, compress the sucrose-versus-plain code, and promote sucrose acceptance.

To directly test whether the reduced discriminative capacity of Earmuff neurons drives sucrose acceptance in the African and Mexican strains, we selectively knocked down *pum* expression in Earmuff neurons of the laboratory strain. Overexpressing *pum-RNAi* specifically in Earmuff indeed led to increased sucrose acceptance (reduced sucrose rejection) in lab strain (**Fig. 5D**). In keeping with the behavioral phenotype, overexpressing *pum-RNAi* in Earmuff neurons increased their response to the plain substrate and hence the plain-to-sucrose response ratio (**Fig. 5E & Extended Data Fig. 6**).

Given Earmuff neurons *increased* their plain response when *pum* was reduced, we were curious if increased *paralytic* (*para*), encoding a voltage-gated sodium channel known to be repressed by *pum* (*21, 27*), might play a role. We overexpressed *para* specifically in Earmuff neurons of the lab strain and found that it indeed increased both plain-to-sucrose response ratio of Earmuff neurons and sucrose acceptance for egg-laying (**Fig. 5F, G, & Extended Data Fig. 6**). Curiously, while overexpressing *pum-RNAi* reduced discrimination by selectively increasing Earmuff’s plain response, overexpressing *para* reduced discrimination by eliciting a larger increase in the plain response and a moderate increase in the sucrose response (**Fig. 5F, G, & Extended Data Fig. 6**).

To further test the idea that *pum* down-regulation – and the consequent *para* elevation – enhances Earmuff neurons’ responses to the plain substrate, we examined these neurons in lab strain males (**Fig. 5H, I**). This is because we found *pum* expression to be sexually dimorphic in the lab strain, with higher levels in male than in female brains (**Fig. 5J**). Thus, Earmuff neurons in lab strain males allows us to assess substrate responses where the *pum* level is naturally elevated. Strikingly, these neurons discriminate sucrose from plain substrates more clearly: compared with females, they responded robustly to sucrose substrates (albeit at a lower magnitude) but virtually not at all to plain substrates, thereby lowering the plain-to-sucrose response ratio (**Fig. 5K**). Importantly, we were able to rescue plain responses in male Earmuff neurons by either selectively reducing *pum* or increasing *para* expression in them, or by examining them in the African strain (surrogate) background (**Fig. 5L**). Thus, our collective results support the notion that reducing *pum* in Earmuff enhances their plain responses, likely by derepressing *para*-mediated excitability. In African and Mexican females, the resulting compressed sucrose-versus-plain response difference drives their acceptance of sucrose for egg-laying (**Fig. 5M**).

## Discussion

In this study, we establish a causal link between naturally occurring genetic variation and strain-specific innate decision bias in *Drosophila*. Our data trace a path from intronic SNPs in the translational repressor *pum*, through value coding in a pair of decision-critical inhibitory neurons, to the resulting behavioral choice. We propose (**Fig. 5M**) that intronic variants in a wild-caught African strain – and in a Mexican strain – lower *pum* expression, thereby derepressing the voltage-gated sodium channel *para* in Earmuff neurons, which directly inhibit the decision center. The ensuing increase in excitability compresses Earmuff’s value code for sucrose versus plain substrates. Furthermore, *pum* knockdown or *para* overexpression specifically in Earmuff neurons of a lab strain recapitulates the wild-caught phenotype, demonstrating that tuning Earmuff activity alone is sufficient to redirect choice. Together, these findings provide an end-to-end example of how natural regulatory variation reshapes circuit properties to alter behavior and a mechanism that may be broadly applicable given the evolutionary conservation of *pum* and *para*.

Is reduced *pum* expression – or elevated *para* – specifically in Earmuff neurons the sole cause of the altered sucrose preference in the two wild-caught strains? We think not. Our PLINK analysis revealed several significant SNPs located in other genes (**Supplementary Table 1**), although their roles remain untested. Further, while we focused on Earmuff neurons here, additional decision-relevant nodes may be affected. Notably, we observed a similar impact of *pum* and *para* on TPN2 neurons, which are presynaptic to Earmuff (**Extended Data Fig. 7**). OviENs, the two pairs of neurons reciprocally connected to oviENs (*12*) (**Fig. 5M**), and oviDNs might all be impacted. Assessing their response properties in laboratory versus wild-caught backgrounds will be a logical next step.

How could *pum* reduction selectively enhance the plain response in Earmuff neurons? One possibility is that Pum proteins are preferentially enriched at synapses receiving the plain-signaling input relative to the sucrose-signaling one. As such lowering *pum* would boost sodium channel locally, selectively elevating response to plain stimuli. Testing this hypothesis will require genetically accessing relevant presynaptic partners of Earmuff neurons and visualizing Pum-dependent translational repression at subcellular resolution. The sexually dimorphic Earmuff responses offer a tractable pilot: in males, Earmuff neurons’ plain response is much reduced compared to in females, hinting enrichment of Pum on synapses receiving plain input.

Finally, could down-regulation of *pum* be a general route to increased sucrose acceptance over plain for oviposition in nature? We believe so. First, line 42 is not the only wild-caught strain failed to complement line 51 (**Extended Data Fig. 2**), implying *pum* might be reduced in those strains as well. Second, the key variants (SNPs) driving sucrose acceptance in line 128 (caught in Maine) must differ from those in line 51, as the two strains complemented each other (**Fig. 1**), yet *pum* expression in 128 was also reduced (**Extended Data Fig. 2**). Thus, variants outside the *pum* locus can still converge on lowering *pum* expression. Adjusting *pum* levels, through diverse molecular mechanisms, may therefore be a common evolutionary lever for tuning sucrose valuation for oviposition. What selection pressures might tip the lever? Possibilities include limited sugar availability and reduced predation risk on sugary substrates in particular environments.

## Methods

### Fly husbandry

Flies were reared on molasses-based food and maintained in a 25°C incubator with 60% humidity. For most behavior and imaging experiments, we used mated females that were 6-14 day old. For experiments where we used RNAi to knock down gene expression, flies were aged up to 22 days to allow sufficient time for targeted mRNA to be depleted. See the table of (abbreviated) crosses used for the flies used for each experiment.

### Immunostaining

Brains and ventral nerve cords (VNCs) were dissected and fixed in 4% paraformaldehyde prepared in PBS containing 0.3% Triton X-100 (PBST). The dilution factors for the antibodies used in this study were as follows: NC82: 1:50, anti-GFP: 1:1000, anti-RFP: 1:5000, anti-V5: 1:500, donkey anti-mouse Cy3: 1:500, donkey anti-mouse Alexa Fluor 488: 1:500, donkey anti-rabbit Alexa Fluor 594: 1:500, and donkey-anti rabbit Alexa Fluor 488: 1:500. Samples were imaged using a Zeiss LSM700 with either a 20x air objective or a 40x oil objective and the acquired confocal images were then post-processed with Fiji.

### RT-qPCR

<u>RNA Extraction</u>: Female fly brains (n = 30) or heads (n = 15) were dissected in RNase-free PBS and transferred to TRIzol reagent. RNA was extracted using the Qiagen RNeasy Kit and Qiagen Shredder column, following the manufacturer’s protocol. <u>DNase treatment</u>: To remove genomic DNA contamination, samples were treated with TURBO DNase™. RNA concentration was measured using the Qubit RNA High Sensitivity Assay Kit. <u>cDNA Synthesis</u>: cDNA was synthesized from total RNA using the SuperScript™ IV First-Strand Synthesis System, following the manufacturer’s instructions. <u>Reverse transcription quantitative PCR (RT-</u> <u>qPCR)</u>: We used QuantStudio3 to perform qPCR using TaqMan™ Fast Advanced Master Mix, following the manufacturer’s protocol. Gene expression levels were assessed using TaqMan Gene Expression Assays with Dm02361909_s1 (act5C, housekeeping gene) and Dm02135527_m1 (*pum*, gene of interest.0

### Full genome sequencing SNP analysis

<u>DNA extraction</u>: We extracted DNA from individual F2 recombinants using a standard protocol(*28*). <u>Whole</u> <u>genome sequencing</u>: We used SNPsaurus and Novogene to perform whole genome sequencing of a total of 192 of the selected F2 recombinants. SNPsaurus and Novogene sequenced 96 flies each using Illumina NovaSeq sequencers, with paired-end reads of 159 bp and 150 bp, respectively, and coverages of 10x and 20x, respectively. <u>Variant calling pipeline</u>. We implemented our own GATK-based variant (SNP and indel) calling pipeline to turn the raw reads (FASTQ files) into a VCF file that gives the variant calls for each fly. The pipeline used GATK tools when available, but we preferred fastp to MarkIlluminaAdapters to take advantage of fastp’s adapter auto-detection. We also wrote a Python script to run pipelines in parallel, making efficient use of modern CPUs’ many cores. For both the pipeline and the script, we strove for low SSD wear. Our pipeline produced by-sample GVCF files, which we combined into a single “cohort” VCF file for all 192 flies. We filtered out genotype calls in the VCF file with genotype quality (GQ) below 13, corresponding to 5% call error probability. <u>GWAS using PLINK</u>. The goal of our GWAS using PLINK(*23*) was to identify variants associated with sucrose acceptance. We used the flies’ preference index (PI) both as quantitative trait and to group flies into “sucrose accepting” and “sucrose rejecting” flies (case/control phenotype). For the group-based analysis, we ran the analysis twice with different thresholds – first for groups “PI≥-0.1” and “PI≤-0.75” (with 49 and 82 flies, respectively) and then for groups “PI≥0” and “PI≤-0.80” (with 38 and 73 flies, respectively). We used PLINK’s max(T) permutation procedure with 5 million permutations and Fisher’s exact test to calculate multiple-comparison-corrected empirical p-values (EMP2). We next performed a quantitative trait analysis using PI as a continuous variable. From this analysis, we specifically selected the genomic control-corrected p-values (GC) and the Bonferroni-corrected p-values (BONF). **Supplementary Table 1** shows the 19 variants from the cohort VCF file that were significantly associated (p≤0.05) with sucrose acceptance or PI for each of the 4 p-values – EMP2 (one for each grouping), GC, and BONF. To identify genes potentially affected by each of the 19 variants, we used SnpEff, and the rightmost column in **Supplementary Table 1** shows those genes and the relative location of the variant.

### Egg-Laying Assay (standard)

<u>Preparation of flies to be used for assay</u>: We used our standard “egg-laying deprivation” protocol previously described(*29*). Briefly, we grouped 35 females with ∼10-15 males in a food vial supplemented with yeast paste (live yeast granules mixed with 0.5% propionic acid) for 4 days. This was to induce females to produce and maintain many eggs in an environment that lacks a good surface for egg-laying due to excess of larvae by day 4. As such, females will readily lay eggs when placed in our custom chambers (apparatuses). <u>Preparation</u> <u>of egg-laying substrates</u>: We prepared the substrates right before dispensing individual females (deprived following the protocol we just stated) into our custom chambers. The sucrose substrates were made by diluting an aliquot of 2M stock solution into 1% agarose (e.g., 750 μl into 10 ml of 1% agarose) while the plain substrates were made diluting the same amount of ddH2O into 1% agarose. We routinely keep a bottle of pre-melted 1% agarose in a 55 °C water bath. After being loaded into the chambers, flies are covered and incubated at 25 °C with 60% humidity for egg-laying. <u>Egg counting</u>: After allowing flies to lay eggs for 18–20 hours (“overnight”), we took pictures of the bottom plate (see **Fig. 1**) and then used our (custom-built) automatic egg counter to segment arenas and count the eggs laid in each trough (code available at https://github.com/rcalfredson/Eggsactly). The tool is a web-based application that automatically quantifies eggs in each agarose region from an overhead photograph of a multi-fly egg-laying chamber. After the user uploads an image, a neural network detects the circular hole at the center of each fly enclosure (arena) and uses these holes as anchor points to define segmentation boxes around the two agarose regions at opposite ends of each arena. Each agarose region is then cropped into its own subimage, which is passed to a second neural network trained to detect individual eggs. Finally, the user can review the automatically identified eggs, correct counts if needed, and download the results either as a CSV (with egg counts per agarose region) or as a fully annotated image. <u>Calculation of Preference Index (PI)</u>: We then calculated the preference index (PI) for each animal using the following formula: (# of eggs on sucrose − # of eggs on plain)/(total # of eggs). This index reflects the relative preference for sucrose over plain agarose. Importantly, if a given fly laid fewer than 10 eggs, its PI was omitted for analysis – we adopted the view that flies’ preferences are clearer if they had laid at least 10 eggs.

### Egg-laying assay (optogenetics)

<u>Fly and chamber preparations</u>: Female flies were prepared and their egg-laying PI was calculated as in the non-optogenetics case, but in the optogenetics case vials were kept in darkness and the food was supplemented with 200 mM all-trans-retinal to allow CsChrimson to be activated by light. Additionally, plain agarose substrates were supplemented with 100 mM sorbitol to ensure flies had access to nutrition during the assay. <u>Egg-laying assay</u>: Flies were loaded into arenas under dim lighting conditions and were allowed to lay eggs overnight (>14 hours) in the SkinnerSys setup, a high-throughput closed-loop stimulation platform. The setup included several custom components optimized for tracking and stimulation: 1) SkinnerTrax: Custom software for real-time tracking of multiple flies while delivering light pulses to individual flies based on their behaviors(*30*) (code available at https://github.com/ulrichstern/SkinnerTrax); 2) Modified two-choice arenas with shorter and slightly angled sidewalls; 3) LED stimulation setup: A custom printed circuit board (PCB) equipped with 80 individually controllable red (624 nm) Cree LEDs, with two LEDs per arena for uniform illumination. 4) Apparatus stand: a custom-designed stand held the arenas, cameras (Microsoft LifeCam Cinema), and PCB in fixed positions. A dimmed light pad (Artograph LightPad A920) provided low-intensity backlighting for tracking without activating channelrhodopsins. Cameras were fitted with blue-pass filters (LEE Filters 172 Lagoon Blue) to reduce the pulsing red LEDs’ impact on tracking. <u>Stimulation protocol</u>: Optogenetic stimulation was applied only when SkinnerTrax detected that a fly was positioned over the agarose substrate of interest. The LEDs for that arena were pulsed at 2 Hz (250 ms on/off), providing approximately 6.8 μW/mm² of red light during “LED on.”

### Calcium Imaging

<u>Preparation</u>: Each fly was anesthetized individually on a CO₂ pad, and the head was positioned through an opening in an aluminum foil plate. UV-cured glue was applied to immobilize its wings and thorax, while the eyes and proboscis were also secured. The head was then submerged in 1X standard imaging buffer(*22*) freshly supplemented with 4 mM CaCl₂. A small section of the cuticle was carefully removed, to expose the region of the interest. An additional 100 µL of 1X imaging solution with 4 mM CaCl₂ was applied to facilitate imaging with a water immersion objective. <u>Image acquisition</u>: Imaging was performed at 25°C using a ZEISS LSM700 laser confocal microscope equipped with a 40X water immersion objective,at 128 x 128 pixels and 8 fps. <u>Stimulation protocol for substrate contacts</u>: During imaging, agarose with 150 mM sucrose or plain agarose was presented to the fly’s legs for about 60 seconds using a micromanipulator. Each substrate was presented in alternating order three times with about 30 second of no substrate in between agarose presentations. <u>Stimulation protocol for solution contacts</u>: To acclimate the fly’s legs to moisture, 100 µL of DI water was first applied, ensuring that subsequent response to the sucrose solution would be specifically due to sugar detection by the leg chemosensory receptors rather than a response to moisture. Following this acclimation step, 50 µL of 450 mM of either sucrose or sorbitol was added to the 100 µL of DI water in which the fly’s legs were submerged. <u>Fluorescence quantification and analysis</u>: Raw LSM files were first uploaded to an in-house application for motion correction to improve accuracy of fluorescence measurements. Regions of interest (ROIs) were defined in ImageJ to measure fluorescence intensity. For Earmuff neurons, the ROI was drawn around the soma in each hemisphere, while for TPN2 neurons, the ROI encompassed the arbors within the SEZ region. For oviDN and oviEN neurons, the ROI was drawn in the SMP region of the brain. Baseline fluorescence (F₀) for each substrate was determined from four to ten time points immediately preceding leg contact with the substrate, depending on the experiment. Fluorescence values (F(t)) were then measured, and the change in fluorescence (ΔF/F) was calculated using the formula: ΔF/F = (F(t) - F₀) / F₀. ΔF/F values were calculated over the entire duration of substrate contact and averaged across three trials for each substrate condition.

### Statistics

All statistical tests, except the SNP analysis, were performed using GraphPad Prism 10. Sample sizes are stated in the figure legends. Unless mentioned otherwise, each dot in a swarm plot and each sample represents a biological replicate (an individual fly). For Ca^2+^ imaging, we typically averaged responses from three technical replicates for the same fly to give one sample. Bars and whiskers indicate mean and 95% confidence interval (CI). Similarly, ΔF/F graphs show mean and 95% CI bands (e.g., Fig. 4H). Also, we used unpaired *t*-tests with Welch’s correction to compare two samples, except for 4I where paired t-tests were used, and one-way ANOVA (with Tukey’s multiple comparison test) to compare three or more samples. To assess whether a sample mean was different from 0, we used one-sample t-tests. The following notation is used to indicate significance: ns: not significant, ****: *p* < 0.0001, ***: *p* < 0.001, **: *p* < 0.01, and *: *p* < 0.05.

### Key resource table

| Reagent/Resource | Source | Identifier |
| --- | --- | --- |
| **Antibodies** |  |  |
| NC82 (mouse) | DSHB | AB_2314866 |
| Anti-RFP (rabbit) | ThermoFisher | AB_560939 |
| Donkey anti-mouse, Alexa-488 | ThermoFisher | AB_141607 |
| Donkey anti-rabbit, Alexa-594 | ThermoFisher | AB_141637 |
| Donkey anti-rabbit, Alexa-488 | ThermoFisher | AB_2535792 |
| Donkey anti-mouse, Cy3 | Jackson ImmunoResearch | AB_2340813 |
| Anti-GFP (rabbit) | ThermoFisher | AB_221569 |
| Anti-V5 (mouse) | ThermoFisher | AB_2556564 |
| **Oligonucleotides** |  |  |
| *pumillio* | Thermofisher | Dm02135527_m1 |
| *actin5C* | Thermofisher | Dm02361909_s1 |
| **Chemicals** |  |  |
| Sucrose | Sigma Aldrich | S0389-5KG |
| UltraPure Agarose | Invitrogen | 16500500 |
| All trans Retinal | Sigma Aldrich | R2500-100MG |
| UltraPure™<br>DNase/Rnase-Free<br>Distilled Water | ThermoFisher Scientific | 10977023 |
| Active dry yeast | VWR | 75860-348 |
| Potassium Chloride | Sigma-Aldrich | P9541-1KG |
| SlowFade™ Diamond<br>Antifade Mountant | ThermoFisher | S36963 |
| Rnase A, Dnase and<br>protease-free | ThermoFisher | EN0531 |
| RnaseZap | ThermoFisher | AM9780 |
| 10X PBS | Invitrogen | AM9624 |
| DEPC-Treated Water | ThermoFisher | AM9920 |
| RNAlater | Sigma-Aldrich | R0901 |
| TRIzol Reagent | Invitrogen | 15596026 |
| Ethyl alcohol | Sigma-Aldrich | E7023 |
| 2-Mercaptoethanol | Sigma-Aldrich | M6250 |
| QIAshredder | Qiagen | 79654 |
| Rneasy Mini Kit | Qiagen | 74106 |
| SuperScript™ IV First-Strand Synthesis System | ThermoFisher | 18091050 |
| TaqMan™ Fast Advanced<br>Master Mix for qPCR | ThermoFisher | 4444557 |
| Molasses food | Archon Scientific, Inc. | B70101 |
| Qubit DNA HS Assay kit | Invitrogen | Q32854 |
| Qubit RNA HS Assay kit | Invitrogen | Q32855 |
| TURBO DNA-free Kit | Invitrogen | AM1907 |
| **Fly strains** |  |  |
| *w<sup>1118</sup>* |  |  |
| *TPN2-GAL4* (regular) | Kim et al.(31) |  |
| *TPN2-ss-GAL4* | Gift from Barry Dickson | SS66351 |
| *Earmuff* | Janelia | SS31063 |
| *Tophat* (1) | Janelia | SS39993 |
| *Tophat* (2) | Janelia | SS39932 |
| *Peafowl* | Janelia | SS37792 |
| *Amulet* | Janelia | SS32423 |
| *Sullivan* | Janelia | SS42639 |
| *Bobber* | Janelia | SS29032 |
| *Turner* | Janelia | SS49422 |
| *Kelp* | Janelia | SS43885 |
| *Hound* | Janelia | SS47232 |
| *UAS>STOP>KIR-eGFP* | Watanabe et al.(32) |  |
| *Otd-FLP* | Asahina et al.(33) |  |
| *UAS>stop>CsChrimson<sup>tdt</sup>* | Watanabe et al.(32) |  |
| *Vgat-LexA* | BDSC | BL-84441 |
| *oviDN-ss-GAL4* | Janelia | SS46540 |
| *Gad-LexA* | BDSC | BL-94893 |
| *UAS-Kir2.1* | Baines et al.(34) |  |
| *Empty-ss-GAL4* | BDSC | BL-79603 |
| *UAS-CsCrimson<sup>venus</sup>* | BDSC | BL-55136 |
| *UAS-CsChrimson<sup>tdT</sup> (x)* | Gift from Toshi Hige |  |
| *lexA-CsChrimson<sup>tdt</sup> (x)* | Gift from Toshi Hige |  |
| *TPN2-LexA* | Gift from Barry Dickson | VT-057358-LexA |
| *LexA-FLP* | BDSC | BL-55819 |
| *LexA-CsChrimson<sup>venus</sup>* | BDSC | BL-55139 |
| *GRASP (spGFPs)* | BDSC | BL-64315 |
| *UAS-GcaMP6f* | BDSC | BL-52869 |
| *UAS-GcaMP6s* | BDSC | BL-42746 |
| Oahu, Hawaii (Line 0) | Cornell Stock Center | 14021-0231.00 |
| Mysore, India (Line 06) | Cornell Stock Center | 14021-0231.06 |
| Captain Cook, Hawaii (Line 54) | Cornell Stock Center | 14021-0231.54 |
| Guam (Line 198) | Cornell Stock Center | 14021-0231.198 |
| Kuala Lumpur, Malaysia (Line 04) | Cornell Stock Center | 14021-0231.04 |
| Cape Town, South Africa (Line 51) | Cornell Stock Center | 14021-0231.51 |
| Vinasco, Mexico (Line 42) | Cornell Stock Center | 14021-0231.42 |
| Coffs Harbour, Australia (Line 18) | Cornell Stock Center | 14021-0231.18 |
| Le Reduit, Mauritius (Line 53) | Cornell Stock Center | 14021-0231.53 |
| *pum* allele | BDSC | BL-6782 |
| *pum* allele | BDSC | BL-97221 |
| *UAS-pum-RNAi* | BDSC | BL-38241 |
| *UAS-para* | Piggott et al.(35) |  |
| *UAS-GcaMP8m* | BDSC | BL-92592 |
| *UAS-GcaMP7b* | BDSC | BL-79029 |
| *UAS-TNT* | BDSC | BL-28996 |
| *Gr5a<sup>LexA</sup>* | BDSC | BL-93440 |
| *Gr64f<sup>LexA</sup>* | BDSC | BL-93445 |
| *UAS-syt-eGFP* | BDSC | BL-33065 |
| **Software** |  |  |
| Automatic egg counter | In-house | https://github.com/rcalfredson/Eggsactly |
| SkinnerSys | In-house | https://github.com/ulrichstern/SkinnerTrax |
| Motion-correction app | In-house | Based on NoRMCorre(36) |
| Prism 10 | GraphPad |  |
| ImageJ/Fiji | https://imagej.nih.gov/ij/ |  |

### Table of crosses used to generate the flies in each figure

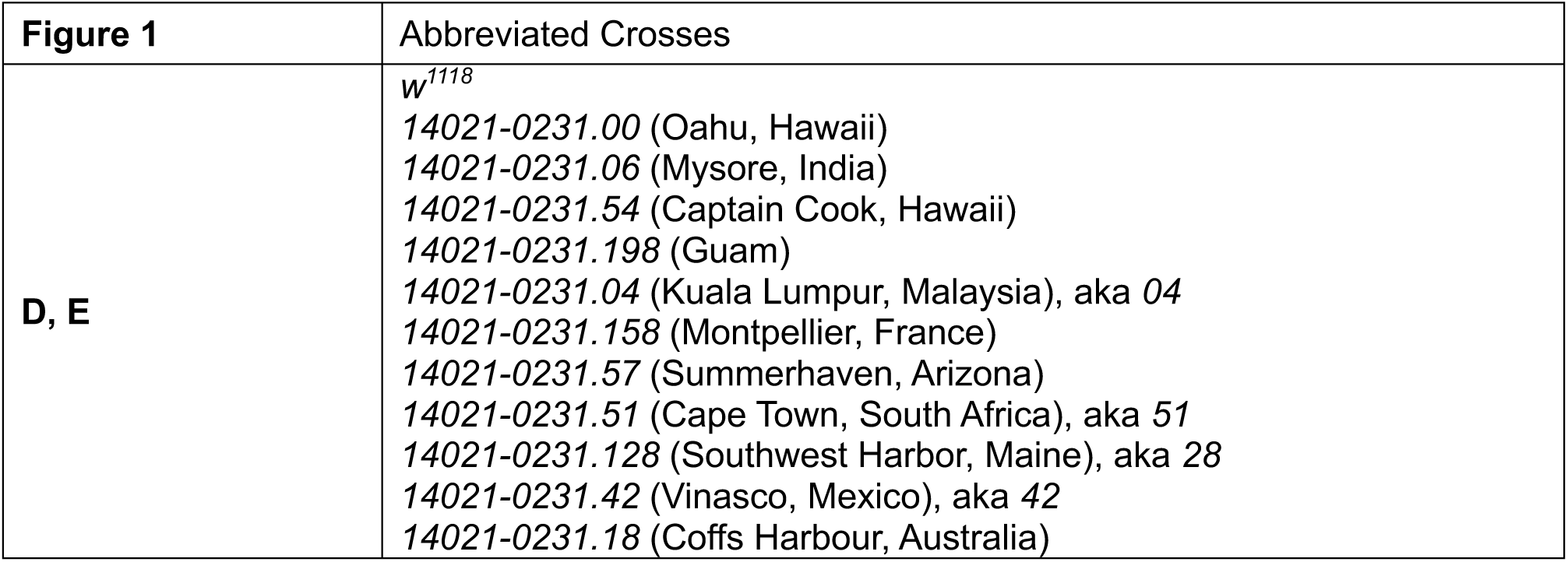
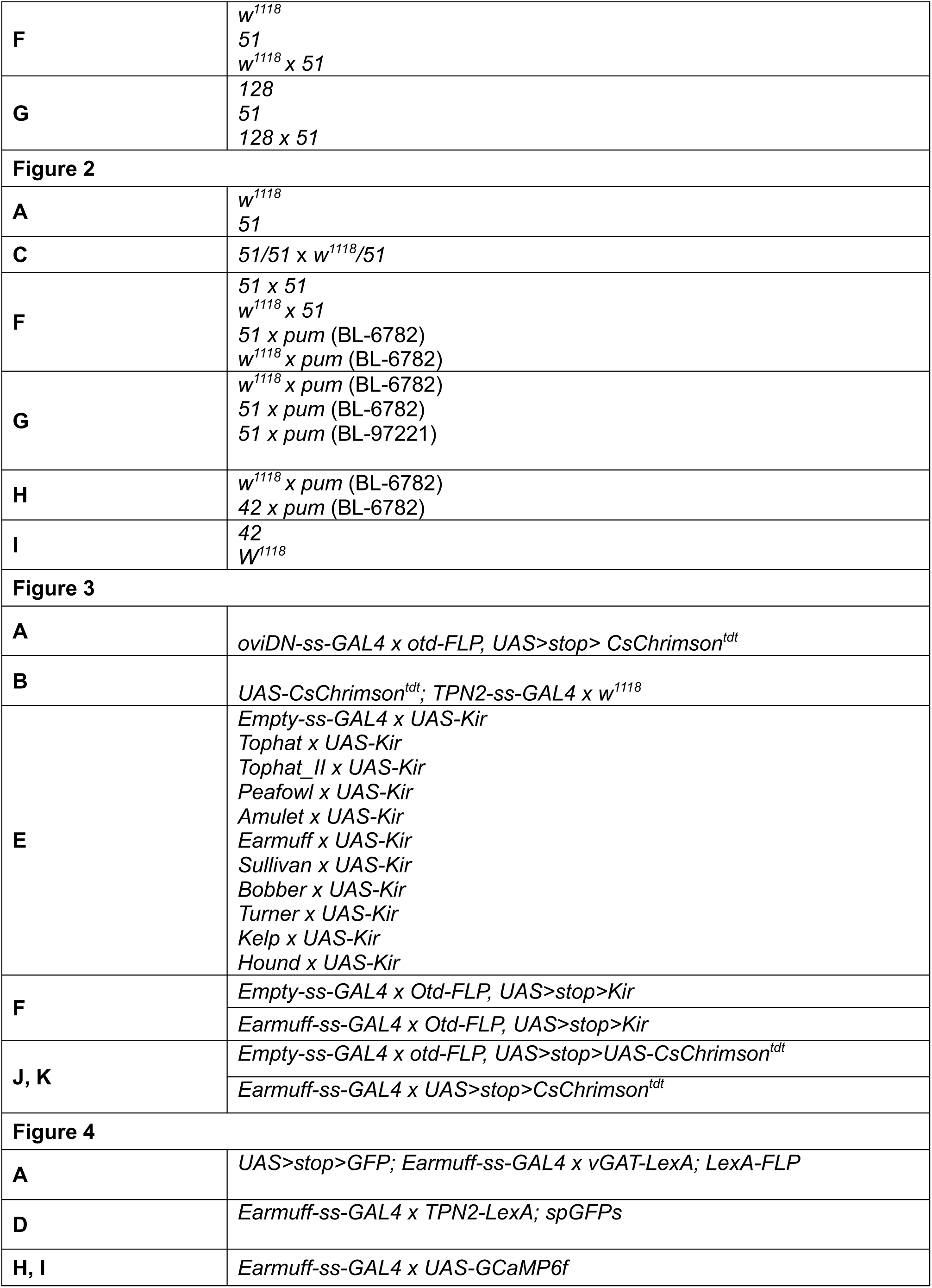
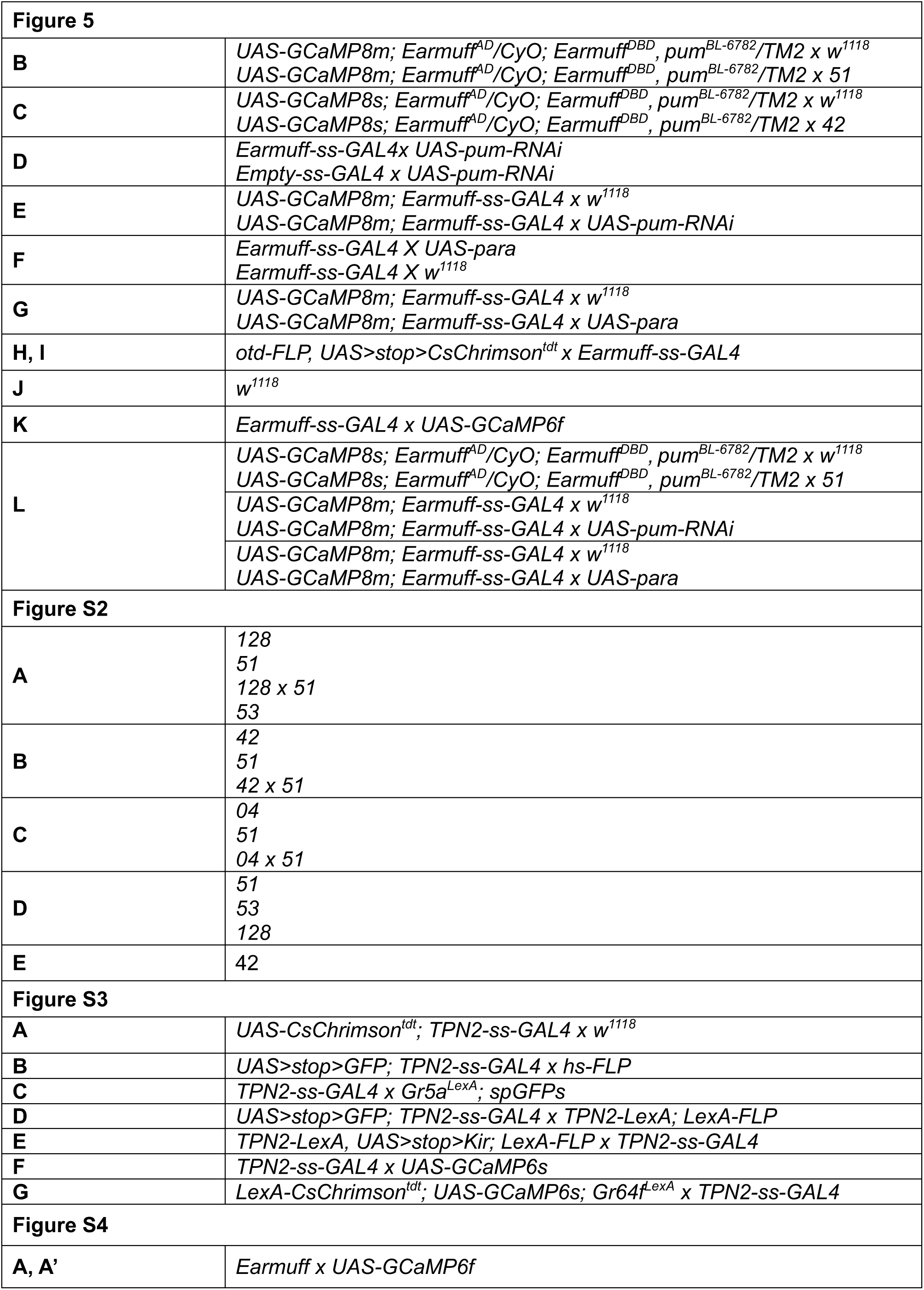
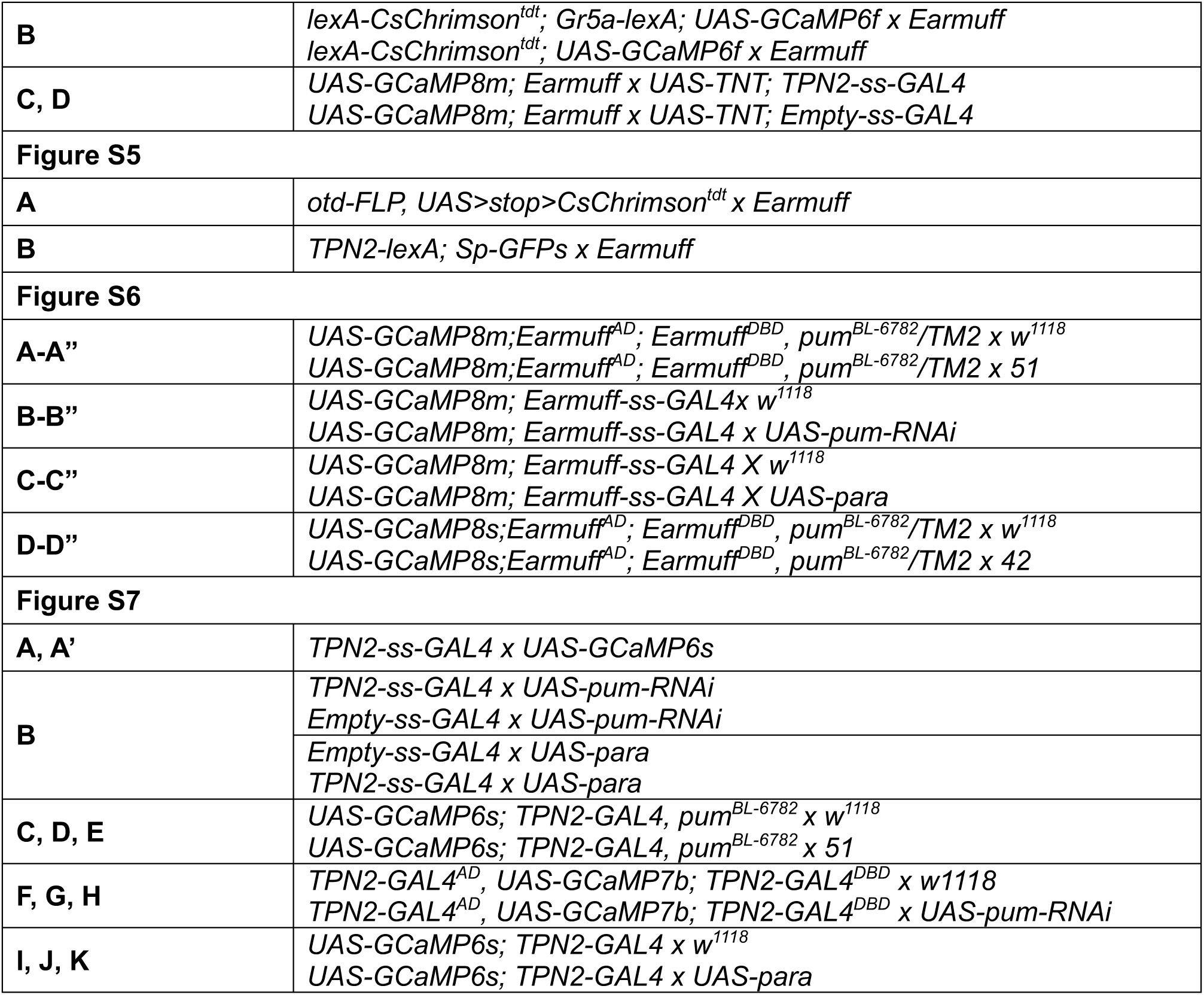

| Figure 1 | Abbreviated Crosses |
| --- | --- |
| **D, E** | $w^{1118}$<br>14021-0231.00 (Oahu, Hawaii)<br>14021-0231.06 (Mysore, India)<br>14021-0231.54 (Captain Cook, Hawaii)<br>14021-0231.198 (Guam)<br>14021-0231.04 (Kuala Lumpur, Malaysia), aka 04<br>14021-0231.158 (Montpellier, France)<br>14021-0231.57 (Summerhaven, Arizona)<br>14021-0231.51 (Cape Town, South Africa), aka 51<br>14021-0231.128 (Southwest Harbor, Maine), aka 28<br>14021-0231.42 (Vinasco, Mexico), aka 42<br>14021-0231.18 (Coffs Harbour, Australia) |
| **F** | $w^{1118}$<br>51<br>$w^{1118} \times 51$ |
| **G** | 128<br>51<br>$128 \times 51$ |
| **Figure 2** |  |
| **A** | $w^{1118}$<br>51 |
| **C** | $51/51 \times w^{1118}/51$ |
| **F** | $51 \times 51$<br>$w^{1118} \times 51$<br>$51 \times pum$ (BL-6782)<br>$w^{1118} \times pum$ (BL-6782) |
| **G** | $w^{1118} \times pum$ (BL-6782)<br>$51 \times pum$ (BL-6782)<br>$51 \times pum$ (BL-97221) |
| **H** | $w^{1118} \times pum$ (BL-6782)<br>$42 \times pum$ (BL-6782) |
| **I** | 42<br>$w^{1118}$ |
| **Figure 3** |  |
| **A** | *oviDN-ss-GAL4 x otd-FLP, UAS>stop> CsChrimson<sup>tdt</sup>* |
| **B** | *UAS-CsChrimson<sup>tdt</sup>; TPN2-ss-GAL4 x w<sup>1118</sup>* |
| **E** | *Empty-ss-GAL4 x UAS-Kir*<br>*Tophat x UAS-Kir*<br>*Tophat_II x UAS-Kir*<br>*Peafowl x UAS-Kir*<br>*Amulet x UAS-Kir*<br>*Earmuff x UAS-Kir*<br>*Sullivan x UAS-Kir*<br>*Bobber x UAS-Kir*<br>*Turner x UAS-Kir*<br>*Kelp x UAS-Kir*<br>*Hound x UAS-Kir* |
| **F** | *Empty-ss-GAL4 x Otd-FLP, UAS>stop>Kir*<br>*Earmuff-ss-GAL4 x Otd-FLP, UAS>stop>Kir* |
| **J, K** | *Empty-ss-GAL4 x otd-FLP, UAS>stop>UAS-CsChrimson<sup>tdt</sup>*<br>*Earmuff-ss-GAL4 x UAS>stop>CsChrimson<sup>tdt</sup>* |
| **Figure 4** |  |
| **A** | *UAS>stop>GFP; Earmuff-ss-GAL4 x vGAT-LexA; LexA-FLP* |
| **D** | *Earmuff-ss-GAL4 x TPN2-LexA; spGFPs* |
| **H, I** | *Earmuff-ss-GAL4 x UAS-GCaMP6f* |

| **Figure 5** |  |
| --- | --- |
| **B** | UAS-GCaMP8m; Earmuff<sup>AD</sup>/CyO; Earmuff<sup>DBD</sup>, pum<sup>BL-6782</sup>/TM2 x w<sup>1118</sup><br>UAS-GCaMP8m; Earmuff<sup>AD</sup>/CyO; Earmuff<sup>DBD</sup>, pum<sup>BL-6782</sup>/TM2 x 51 |
| **C** | UAS-GCaMP8s; Earmuff<sup>AD</sup>/CyO; Earmuff<sup>DBD</sup>, pum<sup>BL-6782</sup>/TM2 x w<sup>1118</sup><br>UAS-GCaMP8s; Earmuff<sup>AD</sup>/CyO; Earmuff<sup>DBD</sup>, pum<sup>BL-6782</sup>/TM2 x 42 |
| **D** | Earmuff-ss-GAL4 x UAS-pum-RNAi<br>Empty-ss-GAL4 x UAS-pum-RNAi |
| **E** | UAS-GCaMP8m; Earmuff-ss-GAL4 x w<sup>1118</sup><br>UAS-GCaMP8m; Earmuff-ss-GAL4 x UAS-pum-RNAi |
| **F** | Earmuff-ss-GAL4 X UAS-para<br>Earmuff-ss-GAL4 X w<sup>1118</sup> |
| **G** | UAS-GCaMP8m; Earmuff-ss-GAL4 x w<sup>1118</sup><br>UAS-GCaMP8m; Earmuff-ss-GAL4 x UAS-para |
| **H, I** | otd-FLP, UAS>stop>CsChrimson<sup>tdt</sup> x Earmuff-ss-GAL4 |
| **J** | w<sup>1118</sup> |
| **K** | Earmuff-ss-GAL4 x UAS-GCaMP6f |
| **L** | UAS-GCaMP8s; Earmuff<sup>AD</sup>/CyO; Earmuff<sup>DBD</sup>, pum<sup>BL-6782</sup>/TM2 x w<sup>1118</sup> |
|  | UAS-GCaMP8s; Earmuff<sup>AD</sup>/CyO; Earmuff<sup>DBD</sup>, pum<sup>BL-6782</sup>/TM2 x 51 |
|  | UAS-GCaMP8m; Earmuff-ss-GAL4 x w<sup>1118</sup> |
|  | UAS-GCaMP8m; Earmuff-ss-GAL4 x UAS-pum-RNAi |
|  | UAS-GCaMP8m; Earmuff-ss-GAL4 x w<sup>1118</sup> |
|  | UAS-GCaMP8m; Earmuff-ss-GAL4 x UAS-para |
| **Figure S2** |  |
| **A** | 128<br>51<br>128 x 51<br>53 |
| **B** | 42<br>51<br>42 x 51 |
| **C** | 04<br>51<br>04 x 51 |
| **D** | 51<br>53<br>128 |
| **E** | 42 |
| **Figure S3** |  |
| **A** | UAS-CsChrimson<sup>tdt</sup>; TPN2-ss-GAL4 x w<sup>1118</sup> |
| **B** | UAS>stop>GFP; TPN2-ss-GAL4 x hs-FLP |
| **C** | TPN2-ss-GAL4 x Gr5a<sup>LexA</sup>; spGFPs |
| **D** | UAS>stop>GFP; TPN2-ss-GAL4 x TPN2-LexA; LexA-FLP |
| **E** | TPN2-LexA, UAS>stop>Kir; LexA-FLP x TPN2-ss-GAL4 |
| **F** | TPN2-ss-GAL4 x UAS-GCaMP6s |
| **G** | LexA-CsChrimson<sup>tdt</sup>; UAS-GCaMP6s; Gr64f<sup>LexA</sup> x TPN2-ss-GAL4 |
| **Figure S4** |  |
| **A, A'** | Earmuff x UAS-GCaMP6f |
| **B** | *lexA-CsChrimson<sup>tdt</sup>; Gr5a-lexA; UAS-GCaMP6f x Earmuff*<br>*lexA-CsChrimson<sup>tdt</sup>; UAS-GCaMP6f x Earmuff* |
| **C, D** | *UAS-GCaMP8m; Earmuff x UAS-TNT; TPN2-ss-GAL4*<br>*UAS-GCaMP8m; Earmuff x UAS-TNT; Empty-ss-GAL4* |
| **Figure S5** |  |
| **A** | *otd-FLP, UAS>stop>CsChrimson<sup>tdt</sup> x Earmuff* |
| **B** | *TPN2-lexA; Sp-GFPs x Earmuff* |
| **Figure S6** |  |
| **A-A''** | *UAS-GCaMP8m;Earmuff<sup>AD</sup>; Earmuff<sup>DBD</sup>, pum<sup>BL-6782</sup>/TM2 x w<sup>1118</sup>*<br>*UAS-GCaMP8m;Earmuff<sup>AD</sup>; Earmuff<sup>DBD</sup>, pum<sup>BL-6782</sup>/TM2 x 51* |
| **B-B''** | *UAS-GCaMP8m; Earmuff-ss-GAL4 x w<sup>1118</sup>*<br>*UAS-GCaMP8m; Earmuff-ss-GAL4 x UAS-pum-RNAi* |
| **C-C''** | *UAS-GCaMP8m; Earmuff-ss-GAL4 X w<sup>1118</sup>*<br>*UAS-GCaMP8m; Earmuff-ss-GAL4 X UAS-para* |
| **D-D''** | *UAS-GCaMP8s;Earmuff<sup>AD</sup>; Earmuff<sup>DBD</sup>, pum<sup>BL-6782</sup>/TM2 x w<sup>1118</sup>*<br>*UAS-GCaMP8s;Earmuff<sup>AD</sup>; Earmuff<sup>DBD</sup>, pum<sup>BL-6782</sup>/TM2 x 42* |
| **Figure S7** |  |
| **A, A'** | *TPN2-ss-GAL4 x UAS-GCaMP6s* |
| **B** | *TPN2-ss-GAL4 x UAS-pum-RNAi* |
|  | *Empty-ss-GAL4 x UAS-pum-RNAi* |
|  | *Empty-ss-GAL4 x UAS-para* |
|  | *TPN2-ss-GAL4 x UAS-para* |
| **C, D, E** | *UAS-GCaMP6s; TPN2-GAL4, pum<sup>BL-6782</sup> x w<sup>1118</sup>*<br>*UAS-GCaMP6s; TPN2-GAL4, pum<sup>BL-6782</sup> x 51* |
| **F, G, H** | *TPN2-GAL4<sup>AD</sup>, UAS-GCaMP7b; TPN2-GAL4<sup>DBD</sup> x w<sup>1118</sup>*<br>*TPN2-GAL4<sup>AD</sup>, UAS-GCaMP7b; TPN2-GAL4<sup>DBD</sup> x UAS-pum-RNAi* |
| **I, J, K** | *UAS-GCaMP6s; TPN2-GAL4 x w<sup>1118</sup>*<br>*UAS-GCaMP6s; TPN2-GAL4 x UAS-para* |

## Acknowledgements

We would like to thank the Bloomington Stock Center for providing fly strains and the Duke Physics Shop for fabricating the behavior apparatuses. We would also like to thank members of the Yang Lab for their comments and suggestions, especially Ned (Suyang) Kan for his help with developing the preparation to image substrate-induced neuronal responses. We would also like to thank Dr. Arnaldo Rosario for his insights on *pum* and Dr. Barry Dickson for providing key reagent and insights on TPN2. This work was supported by an invertebrate neuroscience research grant to C.-H.Y. and by grants from NIH/NIDCD (R01DC018874) and NSF (2034783) to T.H.

## Author contributions

D.M., U.S., and C.-H.Y. designed the project. D.M. performed the bulk of the behavioral and imaging experiments and analyzed the data. S.F., D.M., E.M., Y.C. performed complementation and RT-qPCR experiments and analyzed the data. W.S. and T.H. contributed to the initial Ca^2+^ responses of Earmuff. U.S. designed and built the behavior-dependent stimulation system. U.S. performed the SNP analysis using initial work by R.A. R.A. wrote the code for the automatic egg-counting tool. C.-H.Y. wrote the initial draft of the paper, which was then edited by D.M., E.M., T.H., and U.S.

---

**Extended Data Fig. 1.**
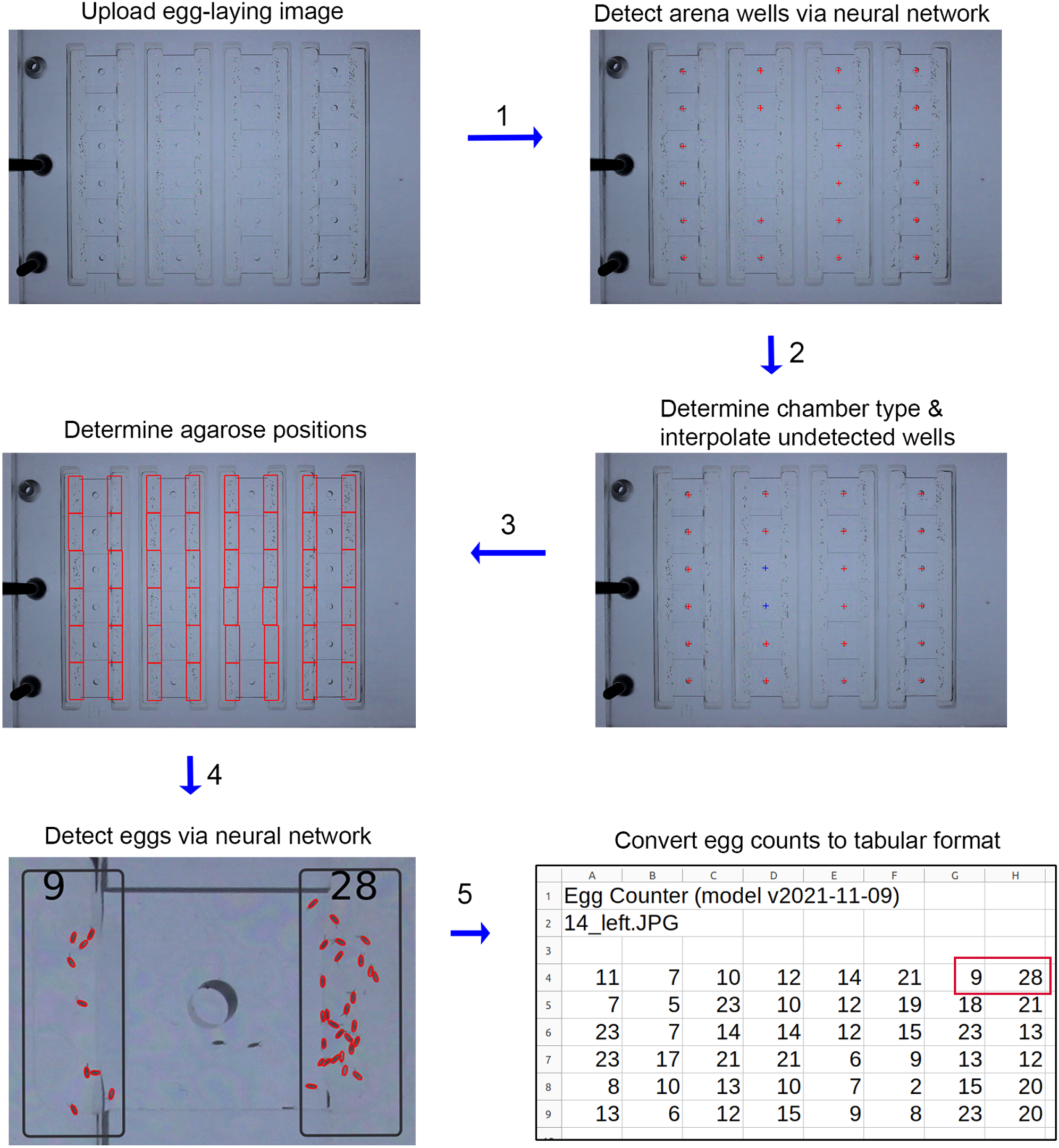
Automated egg-counting workflow. A single script processes a base-plate image and returns egg counts for every arena: **1** Detect the central well in each arena with a convolutional neural network (CNN). **2** Identify chamber type and interpolate any undetected wells. **3** Segment the two agarose substrates per arena. **4** Detect eggs on each substrate with a second CNN. **5** Export counts to a CSV file.

**Extended Data Fig. 2.**
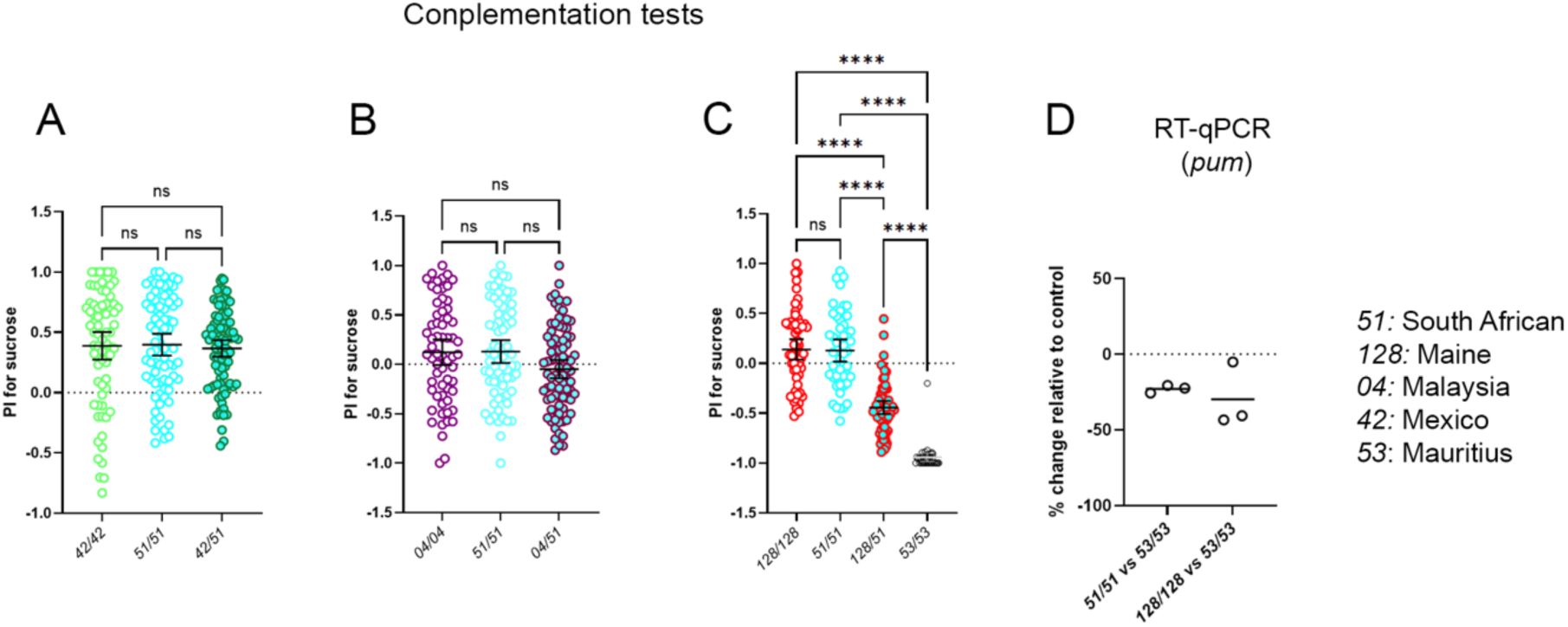
Complementation tests of additional wild strains. **(A**–**C)** PIs of indicated parental lines and their F₁ progeny. **(D)** qRT-PCR showing reduced *pum* expression in the lines 51 and 128 relative to control line 53. Dots represent biological replicates.

**Extended Data Fig. 3.**
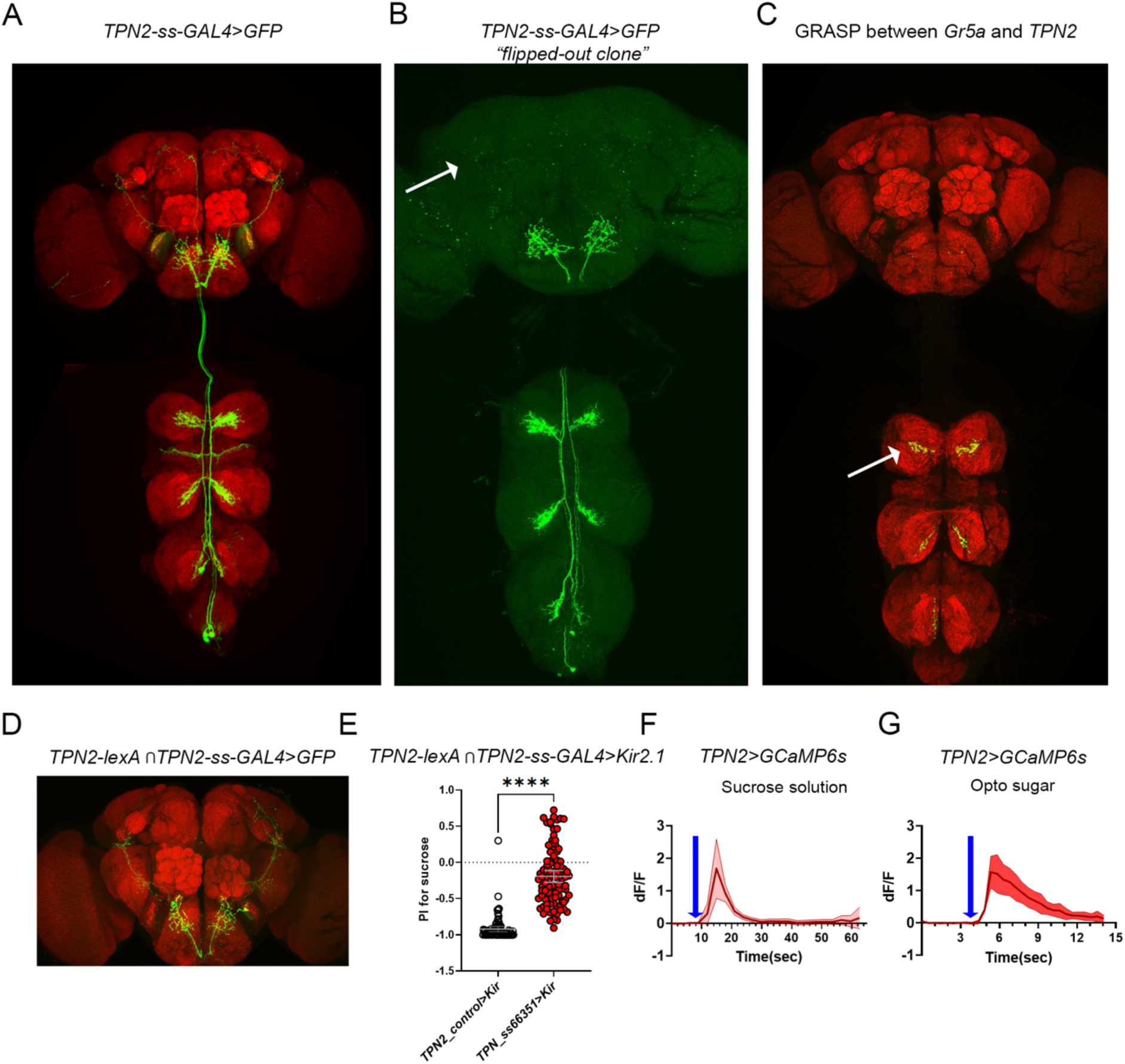
TPN2 neurons are post-synaptic target of leg sweet taste neurons. **(A)** Confocal image of TPN2 neurons labelled by driver *ss66351* (green) with processes terminating both at SLP and SEZ. **(B)** An isolated TPN2 neuron (generated by using random FLP-out) whose axon terminates only at SEZ and not at SLP (arrow). **(C)** GRASP signals between TPN2 and *Gr5a*-expressing sweet neurons in T1 (arrow), T2 and T3 VNC segments. **(D)** Intersectional labelling of *TPN2-LexA* with *ss66351*. **(E)** PIs of controls (n = 85) vs. flies in which the intersected TPN neurons were silenced (n = 96). **(F)** Response of TPN2 to 150 mM sucrose (n = 8); arrow indicates stimulus onset. **(G)** TPN2 responses to optogenetic activation of *Gr64f*-expressing sweet neurons (n = 2 flies, 2–3 technical repeats for each).

**Extended Data Fig. 4.**
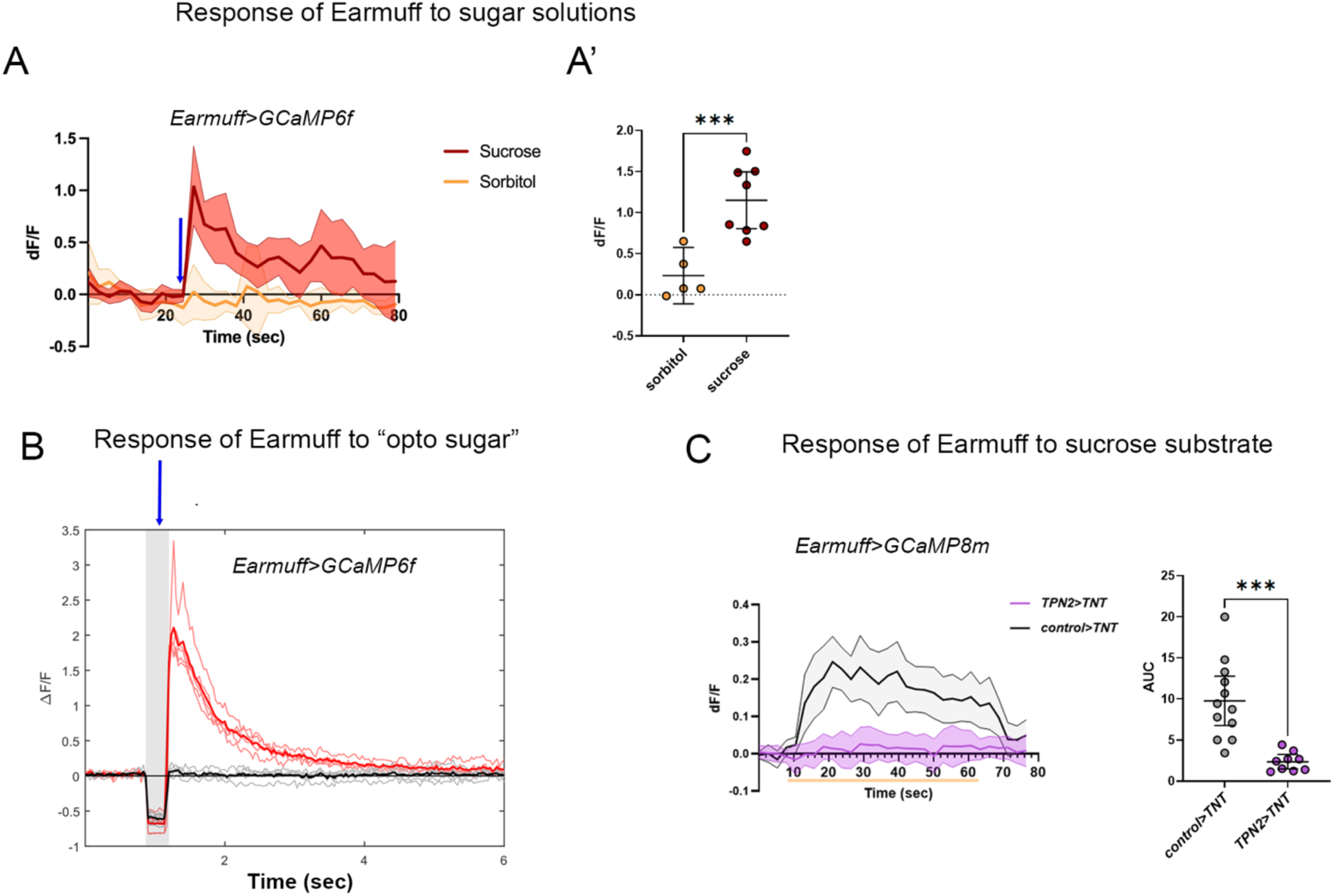
Earmuff sweet-taste responses require functional TPN2. **(A)** Earmuff responses to 150 mM sucrose (n = 8) vs. 150 mM sorbitol (n = 5); arrow, stimulus onset. **(A′)** Peak ΔF/F for flies in (A). **(B)** Optogenetic activation of *Gr5a* (sweet) neurons evokes Earmuff response (n = 5). Shaded area: five 10 ms red-light pulses at 50 Hz; the PMT shutter was closed during stimulation. **(C)** Earmuff responses to sucrose substrate in controls (n = 12) vs. flies with TPN2 synaptic output blocked by TNT (n = 9). Blocking Earmuff itself (*control>TNT*) does not change somatic Ca²⁺ signals.

**Extended Data Fig. 5.**
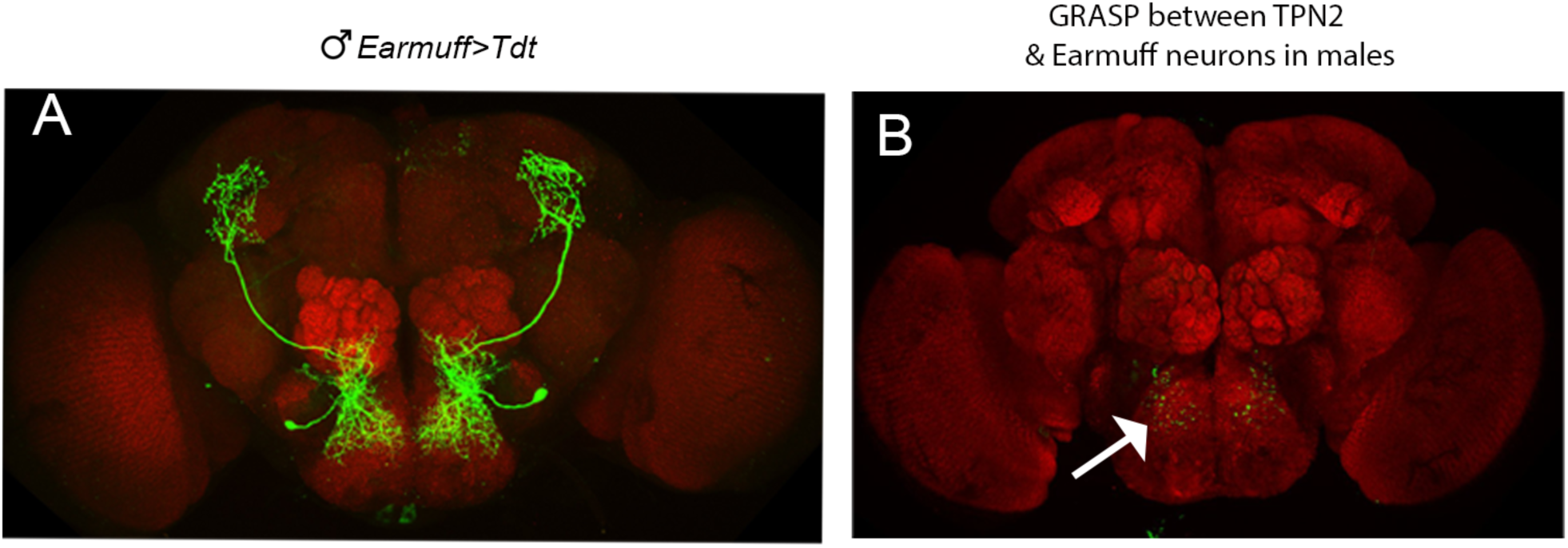
Anatomical Earmuff–TPN2 contacts in male brains. **(A)** Confocal image of male Earmuff neuron morphology (green). **(B)** GRASP between TPN2 and Earmuff neurons in the male SEZ (green puncta and arrow).

**Extended Data Fig. 6.**
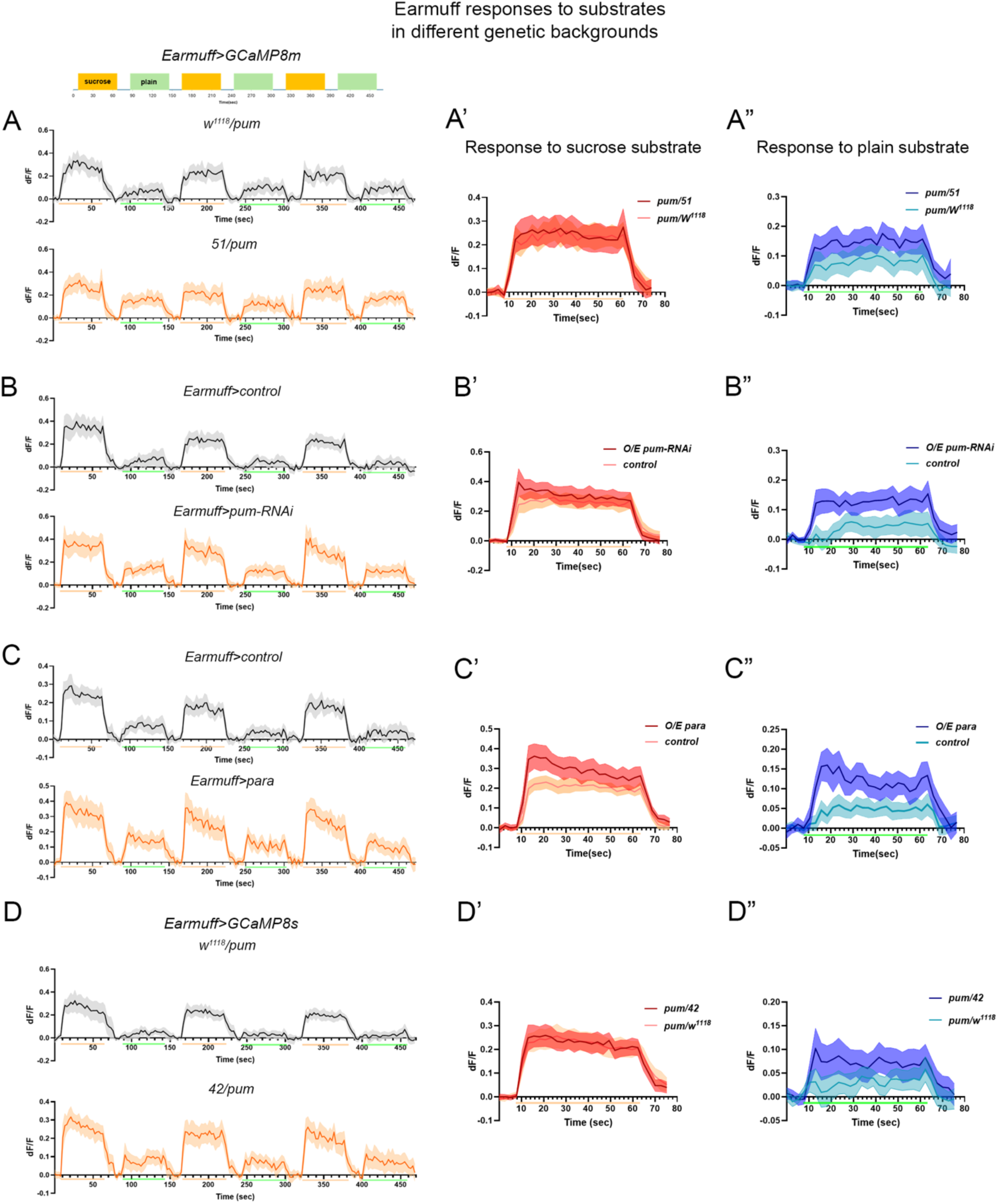
Earmuff responses under different *pum* and *para* manipulations. **(A**, **A’**, and **A”)** Raw traces from and quantification of flies imaged in Fig. 5B during alternating sucrose/plain presentations. **(B**, **B’**, and **B”)** Raw traces from and quantification of flies imaged in Fig. 5E during alternating sucrose/plain presentations. **(C**, **C’**, and **C”)** Raw traces from and quantification of flies imaged in Fig. 5G during alternating sucrose/plain presentations. **(D**, **D’**, and **D”)** Raw traces from and quantification of flies imaged in Fig. 5C during alternating sucrose/plain presentations.

**Extended Data Fig. 7.**
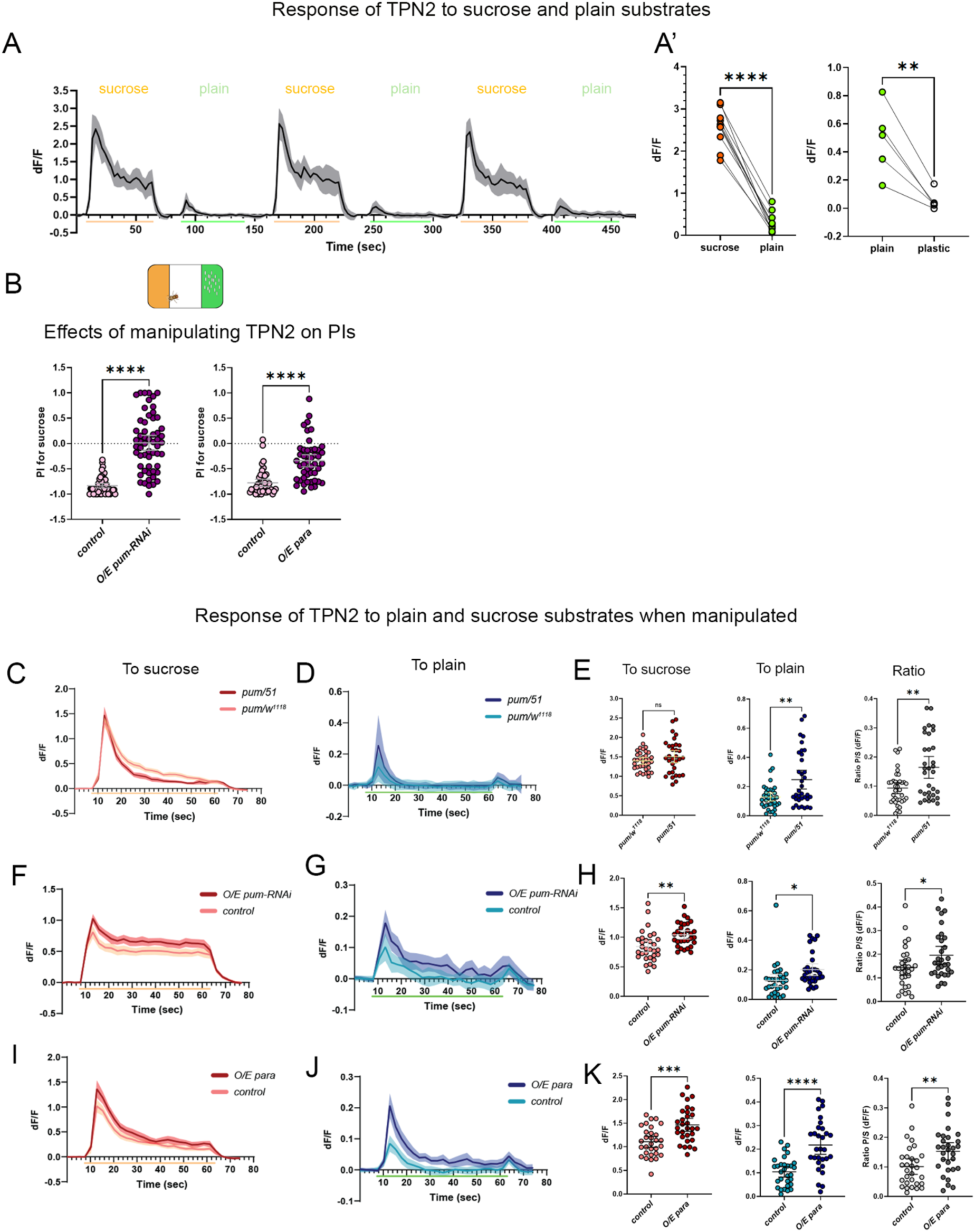
Substrate coding and behavioral impact of *pum* and *para* on TPN2. **(A)** TPN2 responses to sucrose vs. plain substrates (n = 10). **(A′**) Left: max response for sucrose vs. plain (n = 10). Right: max response for plain vs. plastic (n = 5). **(B)** PIs after TPN2-specific *pum* knockdown (n = 57) or *para* overexpression (n = 47) and their respective controls (n = 54, 45). **(C**, **D**, and **E)** Mean TPN2 responses in *pum*/*w*1118 (n = 36) vs. *pum*/51 (African surrogate) flies (n = 32) and quantification of their max responses. **(F**, **G**, and **H)** Mean TPN2 responses in controls (n = 31) vs. *TPN2>pum-RNAi* flies (n = 31) and quantification of their max responses. **(I**, **J**, and **K)** Mean TPN2 responses in controls (n = 30) vs. *TPN2>para* flies (n = 30) and quantification of their max responses.

**Supplementary Table 1.** Additional SNPs and associated genes identified to be significantly associated with PIs.

| Variant |  | Genotype distribution by phenotype group |  | Potentially affected genes |
| --- | --- | --- | --- | --- |
| Position | Seq. change | PI ≥ -0.1 | PI ≤ -0.75 |  |
| 3L:14590265 | T->G | 0/13/25 | 0/56/8 | HGTX (intron) |
| 3L:15778514 | C->A | 0/13/32 | 0/61/17 | lncRNA:CR46197-CG6244 (intergenic_region) |
| 3L:16013248 | G->A | 0/6/30 | 0/48/19 | lncRNA:CR43950 (downstream), lncRNA:CR45998 (upstream), mib1 (upstream), mib1-lncRNA:CR43950 (intergenic_region) |
| 3L:16013253 | GCA->G | 0/6/36 | 0/48/25 | lncRNA:CR43950 (downstream), lncRNA:CR45998 (upstream), mib1 (upstream), mib1-lncRNA:CR43950 (intergenic_region) |
| 3L:16047436 | ATATG->A | 0/11/34 | 0/57/17 | Diap1 (intron,upstream), Mbs (upstream), asRNA:CR45889 (downstream), lncRNA:CR43951 (upstream) |
| 3L:18147732 | A->T | 0/14/33 | 1/59/15 | CG7330 (upstream), geko (downstream), geko-CG7330 (intergenic_region) |
| 3L:18147735 | C->A | 0/14/33 | 1/59/15 | CG7330 (upstream), geko (downstream), geko-CG7330 (intergenic_region) |
| 3L:25627853 | A->AGGTGTTTAC... | 0/13/32 | 0/61/16 | RR48317_transposable_element (upstream), lovit (intron) |
| 3R:8107261 | C->CA | 0/15/28 | 0/61/10 | CG7900 (downstream), FBti0019335 (upstream), puc (intron) |
| 3R:9079556 | C->T | 0/14/23 | 0/60/5 | pum (intron) |
| 3R:11019925 | *->ATGT | 0/12/32 | 0/59/15 | side-VI (intron) |
| 3R:11831160 | TTC->T | 1/16/30 | 0/64/11 | CG6959 (intron) |
| 3R:12966855 | T->C | 0/14/32 | 0/64/15 | CG14384 (intron,upstream), CG7381 (downstream), Cyp304a1 (downstream), Spc25 (downstream), grsm (intron) |
| 3R:13868303 | *->T | 4/11/28 | 30/28/14 | E5 (intron), lncRNA:CR46239 (upstream) |
| 3R:14968498 | T->G | 0/14/31 | 0/60/13 | FBti0062141-lncRNA:CR45599 (intergenic_region) |
| 3R:15125530 | T->G | 0/14/33 | 1/58/14 | CG42788 (intron) |
| 3R:15783287 | *->ACATGTG | 4/11/22 | 31/27/9 | CG5614 (downstream), CG5614-CG9590 (intergenic_region), CG9590 (upstream) |
| 3R:18634603 | CAAAAAAAAA...->C | 0/6/34 | 0/44/18 | CG14301 (intron), qin (intron) |
| 3R:25532557 | TC->* | 3/12/21 | 31/25/12 | msi (intron) |

